# Influence of Reinforcement and Its Omission on Trial-by-Trial Changes of Response Bias in Perceptual Decision-Making

**DOI:** 10.1101/2023.11.02.565253

**Authors:** Maik C. Stüttgen, Andrea Dietl, Vanya V. Stoilova Eckert, Luis de la Cuesta Ferrer, Jan-Hendrik Blanke, Christina Koß, Frank Jäkel

**Author notes:** Correspondence concerning this article should be addressed to Maik C. Stüttgen, Institute of Pathophysiology, University Medical Center of the Johannes Gutenberg University Mainz, 55128 Mainz, Germany.

## Abstract

Discrimination performance in perceptual choice tasks is known to reflect both sensory discriminability and non-sensory response bias. In the framework of signal detection theory (SDT), these aspects of discrimination performance are quantified through separate measures, sensitivity (*d’*) for sensory discriminability and decision criterion (*c*) for response bias. However, it is unknown how response bias (i.e., criterion) changes at the single-trial level as a consequence of reinforcement history. We subjected rats to a two-stimulus two-response conditional discrimination task with auditory stimuli and induced response bias through unequal reinforcement probabilities for the two responses. We compared three SDT-based criterion learning models in their ability to fit experimentally observed fluctuations of response bias on a trial-by-trial level. These models shift the criterion by a fixed step (1) after each reinforced response, or (2) after each non-reinforced response, or (3) after both. We find that all three models fail to capture essential aspects of the data. Prompted by the observation that steady-state criterion values conformed well to a behavioral model of signal detection based on the generalized matching law, we constructed a trial-based version of this model and find that it provides a superior account of response bias fluctuations under changing reinforcement contingencies.

---

Signal detection theory (SDT) constitutes a widely adopted framework for modeling perceptual decisions in psychophysical tasks (for review, see Green & Swets, 1988 and Hautus et al., 2022). SDT breaks down the experimentally observed discrimination performance into two independent performance indices representing sensitivity and response bias. The sensitivity measure *d’* quantifies the degree to which the two stimuli lead to psychophysically discriminable sensations for a given subject. The bias measure (β or criterion *c*) quantifies the degree to which a subject emits one response more frequently than the other. In psychophysics, response bias is usually treated as a nuisance factor, and *d’* is therefore used as bias-free index of perceptual ability. However, the study of bias is interesting in its own right, for example to test some of SDT’s core assumptions such as the shape of the receiver operating characteristic (ROC) curve (Swets, 1961a, 1961b), to investigate mechanisms underlying perceptual learning (Gold & Ding, 2013), and to examine non-sensory factors which influence sensory-guided choices (Alsop, 1998).

There are two well-established procedures to experimentally manipulate response bias in perceptual choice tasks, namely using unequal stimulus presentation probabilities (SPPs; e.g. presenting stimulus 1 more often than stimulus 2) and using unequal payoffs (e.g. providing reinforcement more often for correct choices of one stimulus category than for the other). Neither SPPs nor the payoff matrix are usually made explicit for the subject, but both can be estimated based on the recent history of stimuli, choices, and outcomes. Importantly, SDT itself does not specify how subjects adapt their criterion to a certain experimental situation (criterion learning). Although several models have been proposed as to how subjects may shift their criterion after feedback (e.g., Boneau & Cole, 1967; Busemeyer & Myung, 1992; Dorfman & Biderman, 1971; Erev, 1998; Funamizu, 2021; Kac, 1962; Lak et al., 2017, 2020; Luce, 1963; Mill et al., 2014; Stüttgen et al., 2011; Treisman & Williams, 1984), none of these models has been subjected to extensive experimental scrutiny.

Previous research has shown that a simple income-based criterion learning model is able to fit results from different experiments with rats, pigeons, and mice (Stoilova et al., 2020; Stüttgen et al., 2013; Vandevelde et al., 2023). Here, we test the ability of this model to fit experimental results obtained with rat subjects in two different experiments, and we compare its performance to two related models of criterion learning. In the remainder of the Introduction, we will first explain the concept of adaptive criterion setting within the SDT framework and then introduce the three criterion learning models and describe the design of the two experiments.

## Criterion setting in the signal detection theory framework

We first briefly review how decisions are made within the SDT framework (for a detailed outline, see Hautus et al., 2022). We consider the situation where an observer is performing a two-stimulus two-response conditional discrimination task. This procedure is also referred to as “yes/no”, “single-interval forced choice”, or simply “single-interval” task and should not be confused with the two-alternative forced choice task (Stüttgen, Schwarz, et al., 2011; Wichmann & Jäkel, 2018). SDT posits that each presentation of a stimulus gives rise to a random variable *X* on a decision axis, where *X* is drawn from one of two equal-variance normal distributions which correspond to the two stimuli, S1 and S2 (Figure 1A). The observer decides on the probable identity of the currently perceived stimulus on the basis of a comparison of *X* to a decision criterion *c*. If *X*<*c*, he will respond “S1” (emit R1), if *X*>*c*, he will respond “S2” (emit R2). The distance between the means of the two distributions determines the degree to which a given value of *X* is informative as to the identity of the distribution from which it has been drawn. The distance between the two means divided by their standard deviation is denoted *d’* and constitutes an index pertaining to the discriminability of the two stimuli.

**Figure 1.**
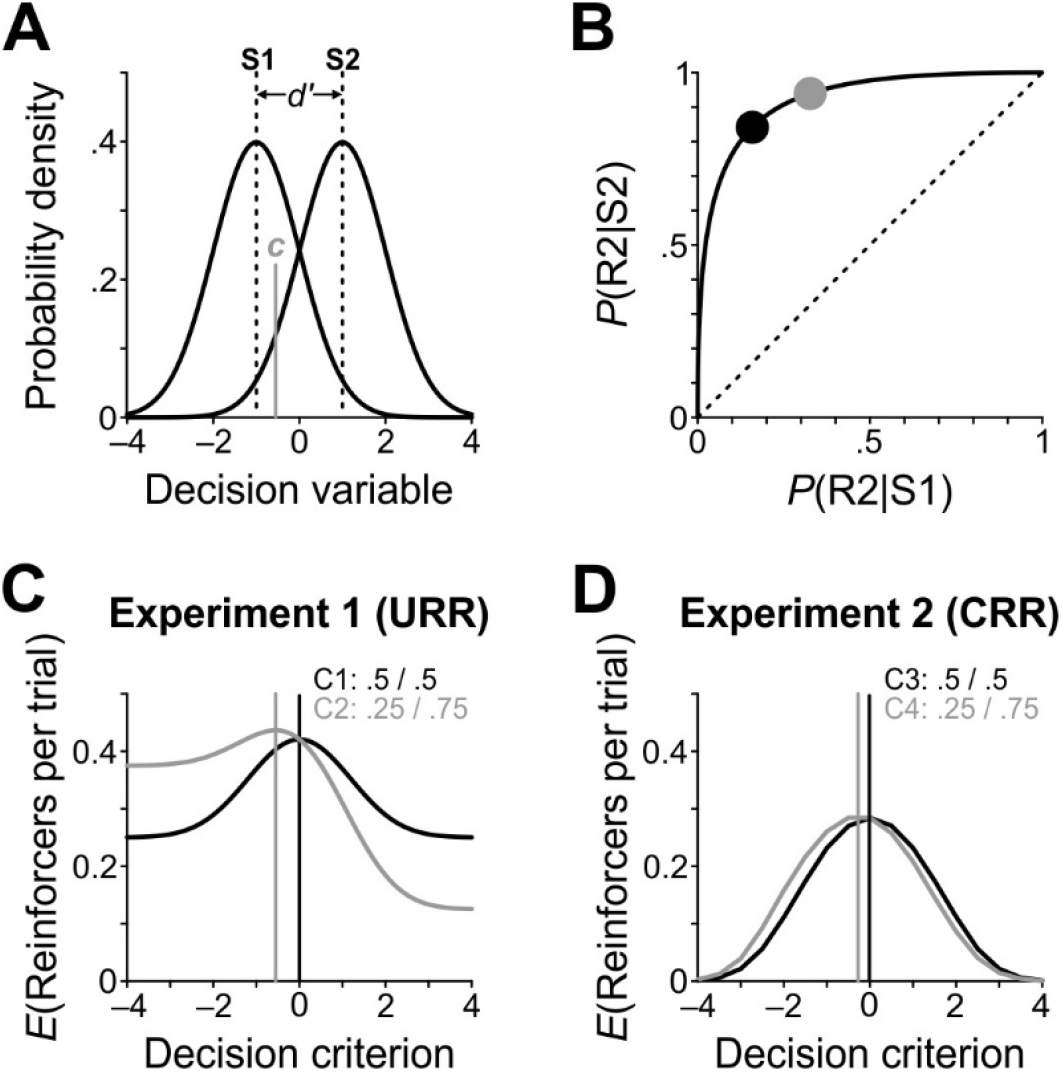
Illustration of criterion setting in Signal Detection Theory. *Note.* A) The two Gaussian distributions correspond to the probability densities of two different stimuli (S1 and S2) on a decision axis (a.k.a. evidence variable). The criterion *c* is located at –0.55 (gray vertical line), thus the observer is biased towards R2. B) ROC curve for *d’*=2. The black dot marks a decision criterion of 0, the gray dot marks *c*=–0.55 as shown in panel A. C) Sample objective reward functions for *d’*=2 and two different sets of reinforcement probabilities (conditions C1 and C2, black and gray curves, respectively) in Experiment 1. Optimal criterion values are represented by vertical lines. D) As in panel C, but for Experiment 2 and conditions C3 and C4.

The actual discrimination performance (i.e., the proportions of correct and incorrect choices in S1 and S2 trials) results from the combination of *d’* and the location of the criterion. If *c* is located exactly halfway between the two means (i.e., if *c*=0), then the percentages of correct S1 and correct S2 trials will be equal (the same is true for incorrect trials). In Figure 1A however, *c*=–0.55, so the observer is biased towards R2. Such a bias can be induced by employing unequal SPPs (here, presenting S2 more often than S1) or by providing reinforcement more frequently in correct R2 than correct R1 trials. Inducing response biases of different magnitudes will yield pairs of hit rates (here, R2 on S2 trials, *P*(R2|S2)) and false alarm rates (here, R2 on S1 trials, *P*(R2|S1)). SDT predicts that these pairs of hit and false alarm rates will all be located on an ROC or “isosensitivity” curve, i.e. a curve containing all possible pairs of hit and false alarm rates for a given *d’* (Figure 1B).

## Three SDT-based models of criterion learning

There is ample evidence suggesting that the criterion is not stationary but affected by stimuli, choices, and outcomes of the immediately preceding trials (e.g., Benjamin et al., 2009; Stoilova et al., 2020; Stüttgen et al., 2011, 2013). The mechanisms underlying this gradual adaptation of the decision criterion are unknown.

Assuming the subject has sufficient experience with the two stimuli to estimate their corresponding distributions, it is reasonable to propose that the criterion is shifted following feedback. In a scenario where correct responses are reinforced and incorrect responses are of no consequence, the simplest criterion learning model would entail shifting the criterion to the left after a correct S2→R2 trial (thus increasing the chance of R2 in the next trial) and vice versa after a correct S1→R1 trial. This mechanism was first proposed by Dorfman and Biderman (1971) but eventually discarded, because the criterion in this model quickly runs off from zero and eventually produces exclusive choice of one response (which happens because of this model’s inherent positive feedback loop). However, this problem can be solved by postulating that the criterion on a trial is an exponentially weighted average of all previous criteria (Stüttgen et al., 2013). In this model (henceforth, “Model 1” or “income-based model”), the criterion updates according to the following equation:

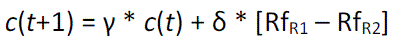

Here, *c*(*t*) is the criterion on trial *t*, γ is a leak factor restricted to range from 0 to 1 which pulls the criterion back towards 0 (the midpoint between the two stimulus distributions) and thus constitutes a leaky integration mechanism, δ is a learning rate parameter which determines the step size of the criterion adjustment, and Rf_R1_ and Rf_R2_ correspond to reinforcement for R1 or R2, respectively, and can take values of either 0 or 1. Thus, the criterion value is incremented by δ if R1 was reinforced and decremented by δ if R2 was reinforced. In trials without reinforcement, both Rf_R1_ and Rf_R2_ are 0, so the criterion is pulled back towards 0 to an extent determined by γ.

However, it is equally conceivable that learning is actually driven by failure to obtain reinforcement (reinforcement omission). In fact, adjusting the criterion (exclusively) on error trials is a mechanism widely believed to hold true in human psychophysics because it predicts the frequently observed phenomenon of probability matching (Dorfman, 1969; Dorfman et al., 1975; Dorfman & Biderman, 1971; Friedman et al., 1968; Killeen et al., 2018; Thomas, 1975). So we conceived of another model (Model 2) which learns exclusively on non-reinforced trials (which not only include all error trials but in our experiment also correct trials in which reinforcement is omitted):

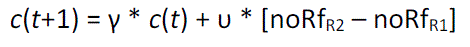

The learning rate parameter for this model is denoted υ (upsilon), and noRf_R2_ and noRf_R1_ represent non-reinforced R2 and R1 trials, respectively, and take a value of 1 when no reinforcement occurs and 0 otherwise. On trials with reinforcement, the term on the right-hand side is 0, thus the criterion is pulled towards 0 at a rate determined by γ.

Finally, Model 3 combines learning after both reinforced and non-reinforced responses:

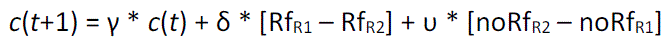

## Description of the experiments

To arbitrate between the three models, we conducted two experiments with rat subjects performing a two-stimulus two-choice conditional discrimination task. Response bias was manipulated by implementing unequal reinforcement probabilities for the two responses, as illustrated in Figure 1. Reinforcement contingencies were unsignaled and changed every five sessions (see Methods for details). The main difference between the experiments were the employed schedules of reinforcement. Experiment 1 featured an uncontrolled reinforcer ratio (URR) schedule in which reinforcements are delivered with a certain probability after correct responses, e.g. *P*(Rf|S1,R1)=P(Rf|S2,R2)=.5, and reinforcements are never delivered after incorrect responses. In this situation, the relative probabilities of obtaining reinforcement for R1 or R2 depend not only on the programmed probabilities, but also on the behavior of the subject. For example, a subject with a criterion of 0 will, in the long run, obtain the same number of reinforcers for both R1 and R2, so the reinforcement ratio Rf_R1_/Rf_R2_ will be 1. However, a subject with a criterion value of –0.55 as in Figure 1AB will emit R2 more often than R1 (in 63% of all trials) and therefore also obtain more reinforcers after R1 than after R2 (in 67% of S1 trials and 94% of S2 trials, so 58% of all reinforcers are produced by R2). So, in this example, the programmed reinforcer probability ratio is 1, but the obtained reinforcer probability ratio is 42/58=0.72.

In Experiment 2, a controlled reinforcer ratio (CRR) schedule was employed. The CRR schedule differs from the URR schedule in the way that reinforcement is allocated to correct responses (see Methods for details). Briefly, the next reinforcement is assigned to either of the two responses with a certain probability (here, .25 for R1 and .75 for R2, a reinforcement ratio of 1:3), and then a variable number of correct responses (VR) of that type have to be completed before that reinforcement is provided, after which the procedure starts over again. The upshot of this procedure is that the reinforcement ratio is fixed, i.e. it does not depend on the response allocation of the animal (McCarthy & Davison, 1984; Stubbs & Pliskoff, 1969). As a result, optimal criterion locations and objective reward functions (ORFs, see Methods for details) differ between experiments even for the same pair of programmed reinforcement probabilities (Figure 1CD).

## Methods

### Subjects

Subjects were four male Long-Evans rats (Janvier Labs), aged eight weeks and weighing 200–250 g at the start of behavioral training. Animals were housed in a common cage in a ventilated temperature– and humidity-controlled cabinet with an inverted day-night cycle (lights off from 8 a.m. until 8 p.m.). Food was available ad libitum in the home cage throughout the entire experiment. Water was freely available on weekends only. During weekdays, access to water was restricted to behavioral testing. The rats’ weight was measured before and after each testing session. Despite water restriction, animals consistently gained weight over the course of the experiments. All procedures were approved by local authorities (Landesuntersuchungsamt Rheinland-Pfalz) and conducted in agreement with German law as well as directive 2010/63/EU of the European Parliament.

### Apparatus and stimuli

Behavioral testing was conducted in a standard operant chamber (ENV-008, Med Associates) measuring 48 cm by 27 cm by 28 cm (LxWxH). The chamber was housed in a sound-attenuating wooden cubicle (length, width, height all 80 cm). One side wall featured three nose ports (LIC.80117RM, Lafayette Instruments) which allowed to detect nose entry and to deliver small amounts of water. Each reinforcement amounted to 30 µl of water which was delivered by 0.5 s of pump activation. House lights provided constant dim illumination but were turned off briefly during time-outs (see below). Experimental hardware was controlled from a PC running custom-written software written in Spike2 via a power 1401 AD converter (Cambridge Electronic Design).

Auditory stimuli were composed of bandpass-filtered white noise bursts with either of two different center frequencies (4096 or 16384 Hz for stimulus 1 and 2, respectively) and durations of 70 ms. Initial training was conducted with easily discriminable stimulus pairs (±0.4 octaves). As rats grew more proficient, bandwidths were gradually increased until animals performed correctly in about 80% of trials with no further improvement (final bandwidths ranged from ±2.8 to ±3.0 octaves, adjusted individually for each subject). White noise was generated at a sampling rate of 200 kHz and filtered in Matlab (The Mathworks). The resulting vectors were imported into Spike2 and output at the same sampling frequency from the analog output port of the AD converter. Sounds were amplified and presented through a loudspeaker attached to the ceiling of the sound-attenuating cubicle. Sound pressure levels of all stimuli were adjusted to 70 dB SPL and calibrated with a ¼ microphone (Microtech Gefell).

### Procedure

Rats were trained on a two-stimulus two-response conditional discrimination procedure (see Supplemental Figure 1A for an outline of a single trial). Rats could initiate trials by poking into the center port continuously for 400 ms. On each trial, one of two stimuli (S1 and S2) was presented, and animals were required to maintain nose poking until stimulus offset. Rats indicated their choice by poking either of the two side ports. A poke into the right choice port (R1) was considered correct following S1, a poke into the left choice port (R2) was considered correct following S2. Correct responses were reinforced according to probabilistic schedules (see below). Correct but non-reinforced responses terminated the trial. A new trial could be initiated immediately. Incorrect responses were punished with a time-out of 4 s during which the house light was turned off. Stimulus sequence was pseudorandomized by concatenating independent sets of 20 trials comprising 10 S1 and 10 S2 trials after shuffling. Within each set, a certain fraction of S1 and S2 trials was assigned reinforcement (if followed by the response designated as correct), corresponding to the reinforcement probability for a given stimulus in a given condition. If rats broke fixation during the 400 ms of trial initiation or during stimulus presentation, the trial was counted as a premature response and aborted. Premature responses were punished with a time-out of 4 s, and aborted trials were not repeated. Each session lasted 45 minutes and contained a median of 551 trials (Experiment 1) and 720 trials (Experiment 2). After 12 weeks of daily training on the task, all rats reliably performed hundreds of trials per day for reinforcement probabilities of .5 (for both stimuli) and performance did not improve anymore.

Reinforcement probabilities used in Experiment 1 were .1, .5 and .9. Four pairs of asymmetric reinforcement probabilities (“conditions”) were presented to each animal, and each condition was in effect for five consecutive days. The sequence of conditions was counterbalanced across animals and is given in Table 1. Before the first experimental condition was run, all animals underwent three days of baseline testing with reinforcement probabilities of both stimuli set to .5. At the conclusion of the experiment, another two days of baseline testing were conducted.

**Table 1.**
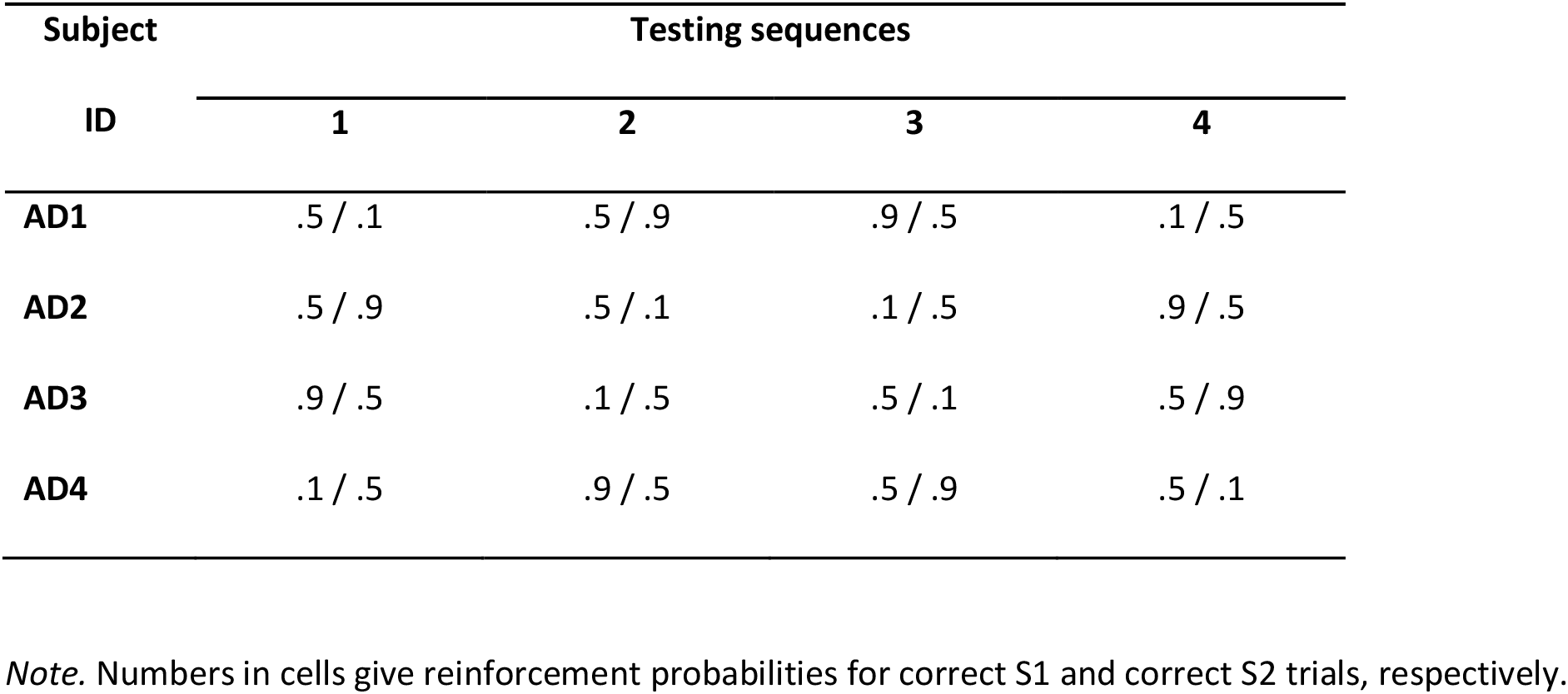
Testing sequences in Experiment 1 (URR) for each of the four animals.

Experiment 2 was similar to Experiment 1, using the same task with the same auditory stimuli, but reinforcement was provided according to a controlled reinforcer ratio (CRR) schedule at intervals determined by two different variable ratio (VR) schedules. Generally speaking, a VR *N* schedule of reinforcement specifies that a reinforcer becomes available after *N* responses. Here, we followed previous authors (McCarthy & Davison, 1984) and implemented a reinforcement schedule where, after each reinforcement, the next reinforcement was assigned to either of the two response ports with a certain probability, and then a variable number of correct responses (VR *N*) toward that port had to be emitted to obtain that reinforcement. Until that happened, no reinforcement could be obtained at either port. This procedure ensures that the relative reinforcement ratio for the two ports is fixed throughout the session. We used relative reinforcement ratios of 1:1, 3:1, and 1:3 (corresponding to relative reinforcement probabilities of .5 vs. .5, .75 vs. .25, and .25 vs. .75). Put differently, as long as the animal emits a minimal number of correct responses in a session, its behavior has no influence on the relative reinforcement ratio. This is in stark contrast to the standard uncontrolled reinforcer ratio schedule, where the animal’s relative response ratio directly influences the relative reinforcement ratio (e.g. the more the animal emits R1, the more reinforcers it will get for R1 and the less for R2).

We ran asymmetric reinforcement conditions with two different VR schedules. Hence, these conditions differ in reinforcement density (maximum number of reinforcers which can be obtained per trial). For VR 2, the number of required correct responses at a given port until reinforcement could be retrieved ranged from 1 to 9 with a mean of 2. For VR 6, the number of correct responses at a given port until reinforcement could be retrieved ranged from 1 to 43 with a mean of 6.

Experiment 2 began with three days of CRR baseline testing where reinforcers became available according to a VR 2 schedule, and reinforcers were assigned to the two ports with equal probability. Then, animals underwent two conditions with VR 2 in which reinforcement probabilities were .25 for S1 and .75 for S2, or vice versa. Next, animals underwent two conditions with the same probabilities but with a lower reinforcement density (VR 6). In each pair of conditions, the order of conditions was counterbalanced across animals and is given in Table 2.

**Table 2.** Testing sequences in Experiment 2 (CRR) for each of the four animals.

| Subject |  | Testing sequences |  |  |
| --- | --- | --- | --- | --- |
| ID | VR 2 |  | VR 6 |  |
|  | 1 | 2 | 3 | 4 |
| **AD1** | .75 / .25 | .25 / .75 | .75 / .25 | .25 / .75 |
| **AD2** | .25 / .75 | .75 / .25 | .25 / .75 | .75 / .25 |
| **AD3** | .75 / .25 | .25 / .75 | .75 / .25 | .25 / .75 |
| **AD4** | .25 / .75 | .75 / .25 | .25 / .75 | .75 / .25 |

*Note.* Numbers in cells specify the relative reinforcement probabilities for correct S1 and correct S2
trials, respectively. All subjects first underwent two conditions with VR 2 and then two conditions with
VR 6.

### Data analysis and model fitting

All analyses and model fits were conducted in Matlab. The experimental data consisted of series of stimulus presentations event time stamps and binary choices. The main indices of behavioral performance were the proportions of correct responses (*P*(correct), calculated as the unweighted mean of *P*(R1|S1) and *P*(R2|S2), and the proportions of R2 (*P*(R2), calculated as the unweighted mean of *P*(R2|S2) and *P*(R2|S1)). Signal detection theory indices were calculated from all completed trials in each session using standard formulae: *d’*=Φ^−1^(HR)– Φ^−1^(FAR), where Φ^−1^ denotes the normal inverse cumulative distribution function, HR denotes the hit rate on S2 trials, *P*(R2|S2), and FAR denotes the false alarm rate on S1 trials, *P*(R2|S1). Relatedly, the criterion was calculated as *c*=–0.5*[Φ^−1^(HR)+Φ^−1^(FAR)]. Optimal criteria (criteria at which expected reinforcement was maximal given a certain value of *d’* and a programmed reinforcement probabilities) were calculated by custom-written Matlab code through numerical optimization.

Details of the fitting procedure for the three SDT-based criterion learning models have been described in earlier studies (Stoilova et al., 2020; Stüttgen et al., 2013). Briefly, the criterion learning models for a fixed leak parameter γ can be formulated as a generalized linear model with a unique maximum, and therefore the likelihood can be maximized reliably with standard numerical optimization procedures (Dorfman, 1973). By repeating this procedure for different γ, the overall likelihood for all parameters can be maximized. The goodness of fit of the three criterion learning models was compared through calculation of the Bayesian Information Criterion (BIC), a dimensionless measure based on the maximum likelihood estimate and controlling for different numbers of parameters in competitor models:

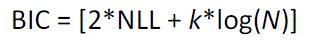

NLL is the negative log likelihood of the data under the model using the best-fitting parameters, *k* is the number of free parameters (four for models 1 and 2, namely γ, the model-specific learning rate parameter (δ or υ) and the two means of the stimulus distributions, and five parameters for Model 3 which features two learning rates), and *N* is the number of trials. Models with smaller BIC values are preferred. As per convention, strength of evidence against the model with the higher BIC value is judged to be “unimportant” if difference in BIC is 0–2, “positive” for differences of 2–6, “strong” for differences of 6–10 strong, and “very strong” if the difference exceeds 10 (Kass & Raftery, 1995).

For Experiment 1, we computed model predictions of the steady-state criterion positions. The responses are stochastic, so there is an equilibrium distribution for the criterion. While deriving this distribution (and showing that it actually exists) is beyond the scope of this paper, heuristically the following can be said: In the steady-state, the expected criterion update is zero. So the expected update due to a reinforced response (for Model 1) / non-reinforced response (for Model 2) is balanced out by the term with the leakage factor that pulls the criterion back to its neutral position.

For Model 1 this means

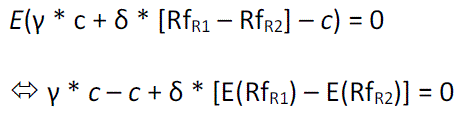

and for Model 2

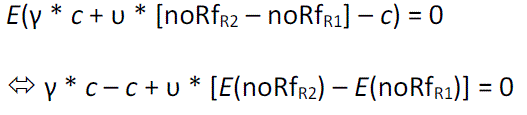

Hence, there is a linear relationship between the criterion position in the steady-state and the difference of the probabilities for (non-)reinforcement for a category 1 response and a category 2 response:

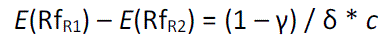

and

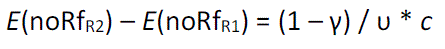

for the two models respectively.

For Experiment 1, the expected values in these equations are straightforward to determine for a given c: In each trial, the expected probability to obtain reinforcement when responding correctly is fixed, so e.g.

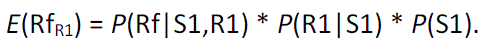

The solutions to these steady-state equations were calculated by custom-written Matlab code through numerical optimization. A graphical example is shown in Supplemental Figure 3.

For Experiment 2, calculating the models’ criterion predictions is not easily possible, because the expected probability to obtain reinforcement when responding correctly depends on the history of stimuli and responses from previous trials, so there are no trial-independent expressions for *E*(Rf_R1_), *E*(Rf_R2_), *E*(noRf_R1_) and *E*(noRf_R2_). We therefore turned to estimate expected criterion values through steady-state criterion values obtained from forward simulations. To that end, each model encountered each of the two experiments 100 times, and expected criteria were obtained as the averages of the 100 criterion values computed over the last 500 trials of each condition (each condition encompassed 1500 trials).

Response ratios and reinforcement ratios for assessing the fit of the Davison-Tustin (DT) model (Davison & Tustin, 1978) was taken from the last two sessions of each condition and were computed separately for S1 and S2 trials. A linear fit was determined for each stimulus type, and the DT sensitivity and bias parameters were determined from the fitted values as described by the original authors:

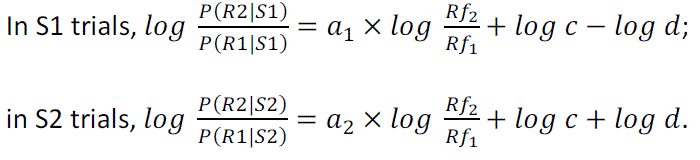

Log *d* represents the vertical separation between the two fitted lines, log *c* represents the overall (stimulus-independent) response bias (not to be confused with the criterion *c* in the signal detection models above), *a*_1_ and *a*_2_ represent the degree to which the response ratio changes when the obtained reinforcement ratio changes. The response probabilities on the left can be estimated from the data. Rf_1_ and Rf_2_ are the reinforcement rates for the R1 and R2 responses, respectively. Reinforcement rates were also estimated from the data; for the CRR schedule, they are by design almost identical to the programmed rates. With these empirical estimates, all the parameters can be found by linear regression (but see (Davison & McCarthy, 1981)). From the resulting regression lines, we can in turn determine the predicted response probabilities of the fits for each condition. In the CRR case, the log odds of the predicted response probabilities are directly given by the regression line. In the URR case, Rf_1_ = *E*(Rf_R1_) = *P*(Rf,S1,R1) = *P*(Rf|S1,R1)\**P*(S1)\**P*(R1|S1), and therefore both sides of the regression equation depend on the response probabilities *P*(R1|S1), and equivalently for Rf_2_. Hence, the predicted response probabilities have to be obtained by solving the fixed-point equation, which can easily be done numerically (e.g. by fixed-point iteration or root finding). The resulting response probabilities can then be used to compute predicted criterion values for a signal detection model.

We further implemented a trial-by-trial version of the DT model. In this model, estimates for Rf_1_ and Rf_2_ in each trial are computed by leaky integration of the reinforcements obtained in past trials, i.e. Rf_1_(*t*) = γ*Rf_1_(*t*–1) + Rf_R1_(*t*) and Rf_2_(*t*) = γ*Rf_2_(*t*–1) + Rf_R2_(*t*), where Rf_R1_(*t*) is 1 if a reward was obtained for an R1 response in trial *t* and 0 otherwise, and analogously for Rf_R2_. Then, log(*P*(R2|S)/*P*(R1|S)) is computed according to the DT model for the stimulus S presented in trial t, and a response is sampled randomly with the probability for an R2 response being *P*(R2|S). For a given γ, the DT model is, again, a generalized linear model with a convex NLL, so the maximum likelihood fit for the model can be found reliably through numerical optimization.

## Results

The left panels in Figure 2 shows *d’* and *c* for all four subjects and for both experiments (see Supplemental Figure 1B for the proportions of correct trials, R2, and aborted trials). Qualitatively, a few observations are obvious: First, asymmetric reinforcement probabilities induced response biases (*c*) in all animals, and these were more pronounced in Experiment 1. Second, discrimination performance (*d’*) did not change as prominently and systematically as response bias, but there seemed to be a trend towards increasing performance over time.

**Figure 2.**
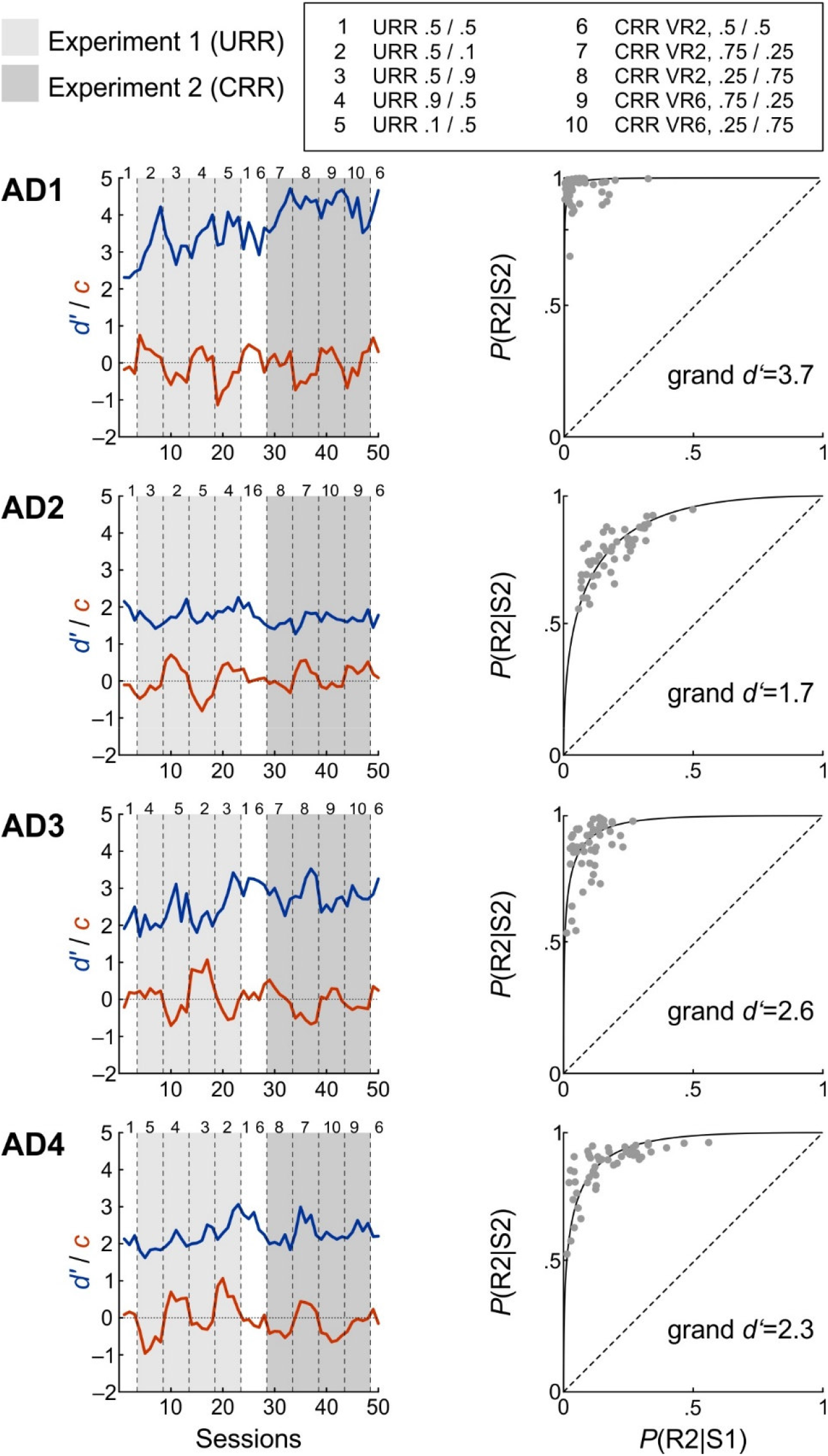
Sensitivity (d’), criterion (c), and ROC plots for all subjects for Experiments 1 and 2. *Note.* Left panels: SDT indices *d’* and *c* for subjects AD1 thru AD4 for both experiments. Sessions belonging to Experiment 1 and Experiment 2 are highlighted by light gray and dark gray shading, respectively. Conditions within experiments are separated by vertical dashed lines in each panel and are referenced by numbers (see inset). Right panels: ROC plots for the four animals. Each data point represents *P*(R2|S2) (ordinate) and *P*(R2|S1) (abscissa) in one of 50 individual sessions. ROC curves were calculated on the basis of the grand *d’* for each animal.

To examine the extent to which *d’* increased over the course of the experiments, we performed linear regression analysis on all *d’* values as a function of session number, separately for each animal. Regression coefficients were statistically significant in three of the four animals (*p*<.02 for AD1, AD3, and AD4 and *p*=.13 for AD2) and positive in three out of four animals (0.034, –0.003, 0.018, and 0.008 for rats AD1 thru AD4, respectively). Thus, despite several months of previous training, three of the four animals exhibited further increments in *d’* over the course of testing, although the increase was small within an individual experiment.

The right panels in Figure 2 show ROC plots for each animal, using data from all experimental sessions. We computed ROC curves for the equal-variance Gaussian model and calculated grand *d’* from averages of *P*(R2|S2) and *P*(R1|S2) of all experimental sessions. Overall, grand *d’* varied from 1.7 (AD2) to 3.7 (AD1), and the distributions of individual sessions’ data points were consistent with a singular curvilinear ROC curve. We thus proceeded to apply SDT-based analyses.

### Comparison of three trial-by-trial criterion learning models

We compared the three criterion learning models in their ability to fit the experimental data. As detailed in Introduction, Model 1 learns exclusively after all reinforced responses, Model 2 learns exclusively after all non-reinforced responses, and Model 3 learns after both. We here report data from fits to the complete data of each subject (i.e., both Experiment 1 and 2) for convenience, since fitting the models separately for the two experiments produced similar results.

Figure 3 shows the fits of the three models to the data of each animal. Qualitatively, the best fit was achieved by Model 3, which was expected given that it is the only model which has two learning parameters. Model 1 was able to capture the general direction of the bias but the fits diverged markedly in some conditions. Finally, Model 2 fared by far worst.

**Figure 3.**
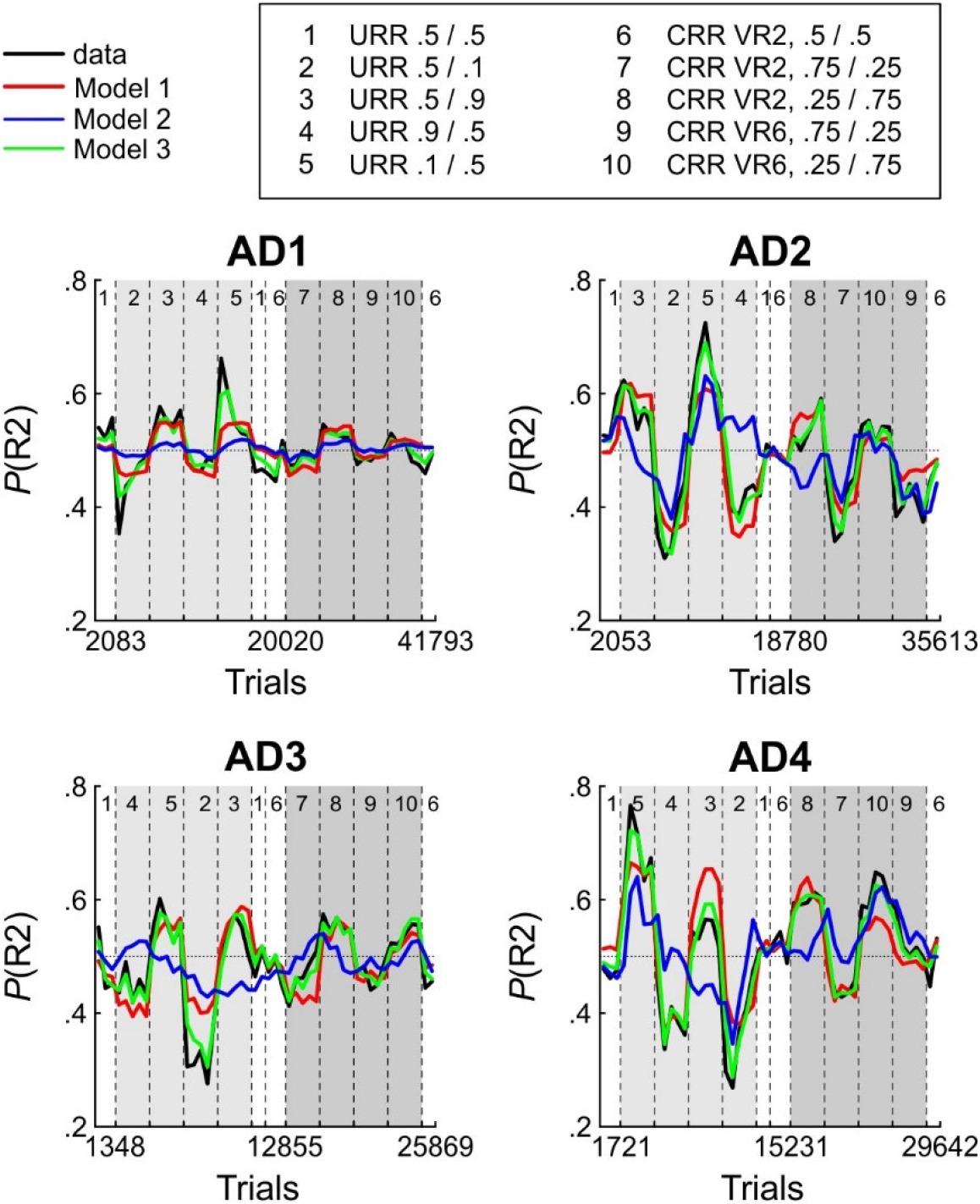
Maximum likelihood fits for the three SDT-based criterion learning models. *Note.* Models are color-coded, empirical *P*(R2) values redrawn in black from Figure 2. All conventions as in Figure 2.

To compare model fits quantitatively, we computed the Bayesian Information Criterion (BIC) for each fit. Figure 4A shows BIC values for each of the four rats’ data fit by each of the three models. Confirming visual inspection, Model 2 was dramatically inferior to Model 1 (BIC differences from 500 to 1000), which again was clearly inferior to Model 3 (BIC differences of 300 to 500), and this was the case for all animals, although all models produced similar estimates of *d’* and γ for a given animal (Figure 4B, left and middle panels). Examination of the learning rate parameters revealed that, while values were always positive for criterion shifts after reinforcement (δ, models 1 and 3), values for criterion shifts after reinforcement omission (υ, models 2 and 3) were, with a single exception, all negative. Importantly, negative learning parameters do not make sense from a theoretical point of view. In the present experimental scenario, they imply that an agent learning from non-reinforced responses shifts his criterion such that he is more likely to produce the same non-reinforced response again in the next trial. In a scenario in which most correct responses are reinforced and incorrect responses are not reinforced, this would increase the likelihood to obtain no reinforcement in the next trial as well, which we confirmed through forward simulations for our experimental conditions (Supplemental Figure 2).

**Figure 4.**
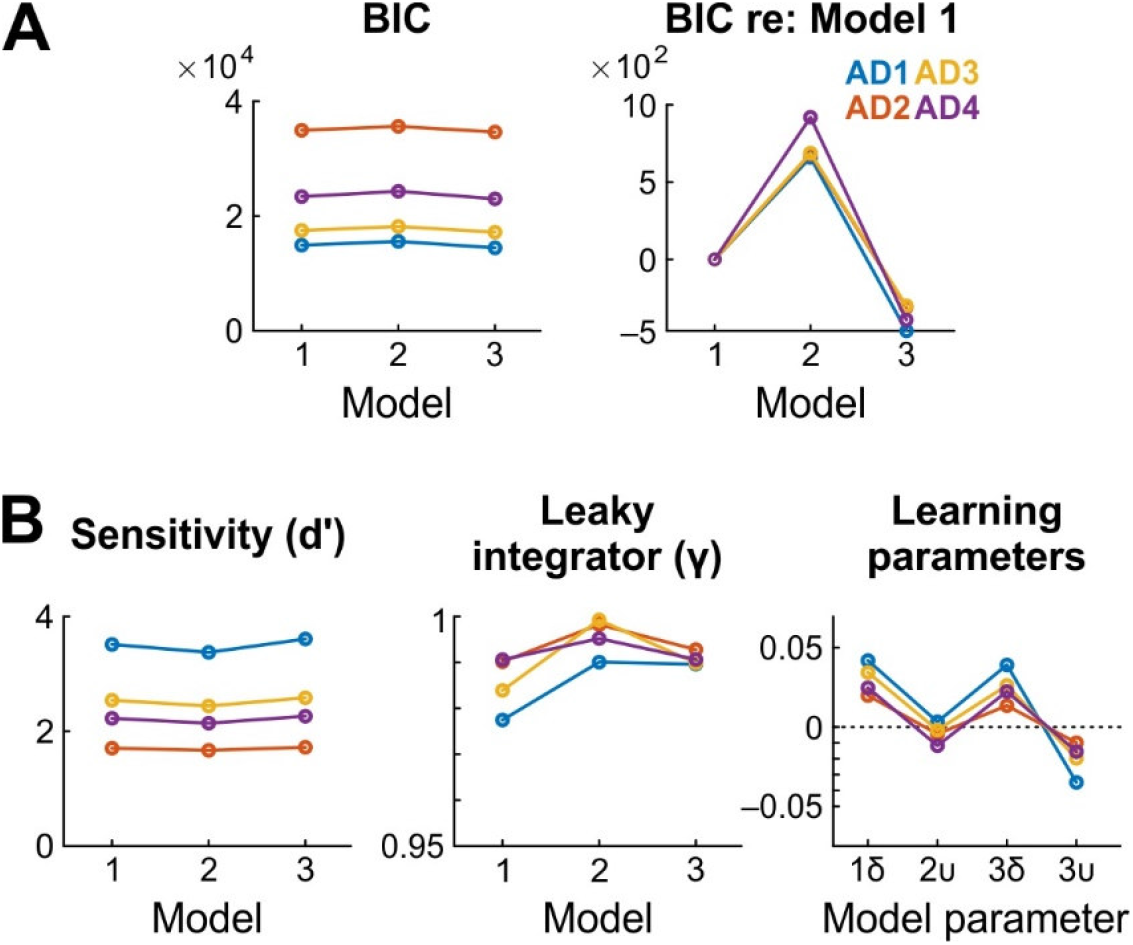
Quantitative comparison of the criterion learning models. *Note.* A) Left panel shows BIC values for each of model for each animal. Right panel shows the same, but normalized to Model 1 for better comparability. The model with the lowest BIC is considered to provide the best fit. B) Left: sensitivity (*d’*) as estimated from each of the three models for each of the four animals. Middle: best-fitting γ values. Right: best-fitting δ and υ values.

### Comparison of predicted and observed steady-state criterion values

In the preceding section, we examined how well the criterion learning models fit the behavioral data on a trial-by-trial basis. The fits were rather bad for Model 1 and Model 2 compared to Model 3, but the negative learning rates for υ in Model 3 do not make sense for theoretical reasons. This analysis, however, does not tell us why Model 1 fails so badly. Another way to evaluate the models is to ask to what extent their predicted steady-state criterion values align with the empirically obtained steady-state criterion values. As explained in Methods and demonstrated in Supplemental Figure 3, the models can be used to predict what value the criterion will converge to for a given a set of model parameters (stimulus means, γ, δ, υ) and reinforcement contingencies. In general, this constitutes an additional and important way to evaluate the models, since a successful model fit does not guarantee that a model endowed with the fitted parameters will generate behavior quantitatively and qualitatively similar to that of the subject if confronted with a different sequence of stimuli and/or reinforcements than those used for the fit (see (Corrado et al., 2005) for an instructive example). Here, we additionally wanted to test whether the bad fits of Model 1 might be explained by its inability to explain the steady-state behavior of the animals.

We therefore generated steady-state predictions for the three criterion learning models for each subject’s fitted parameters and all conditions in both experiments. Furthermore, we compared actual criteria with optimal criteria which provide a convenient benchmark to gauge performance. Assuming that over the course of evolution non-optimal strategies have been eliminated (as they are by definition inferior to optimal ones) may be taken to imply that the brain indeed instantiates an optimal choice algorithm and therefore represents certain quantities such as posterior odds (Harley, 1981; Yang & Shadlen, 2007). In non-human animals, stimulus discrimination performance has been found to come close to optimality in some settings (Feng et al., 2009; Stüttgen, Yildiz, et al., 2011) but not in others (Stoilova et al., 2020; Stüttgen et al., 2013; Teichert & Ferrera, 2010).

Figure 5 shows scatterplots of predicted against actual criterion values for each model. For models 1 and 2 as well as optimality, correlations between predicted and actual values were high and statistically significant for all criterion setting accounts in both experiments (r ranged from .74 to .92, all p-values were <.001). In contrast, predicted and actual criterion models for Model 2 were negatively correlated, as expected based on the negative learning rates obtained from fits of Model 2 and the simulations shown in Supplemental Figure 2, again demonstrating the inadequacy of this model (*r*=–.78 and *r*=–.31 for Experiment 1 and Experiment 2, respectively). We will not consider Model 2 any further.

**Figure 5.**
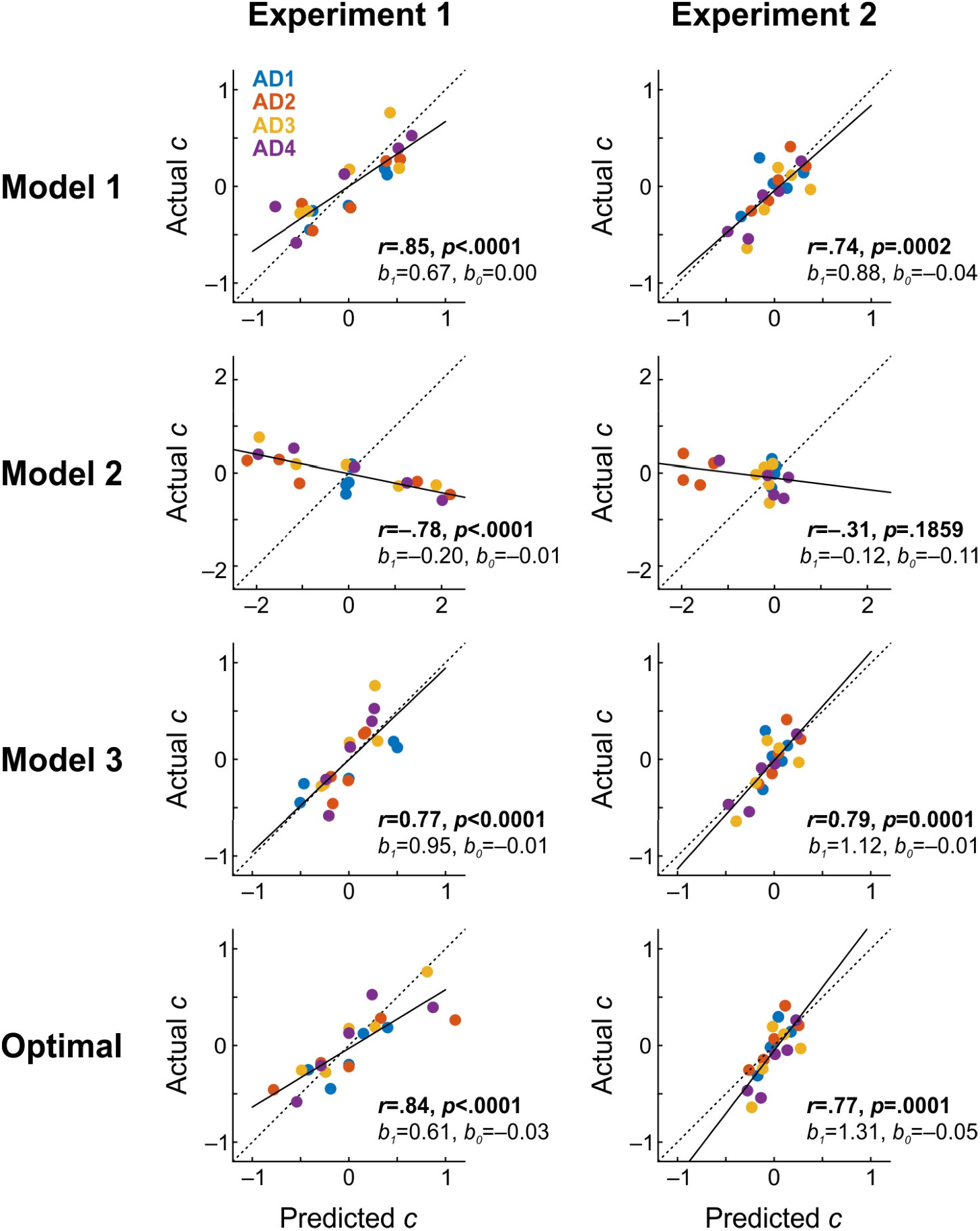
Comparison of predicted and actual steady-state decision criteria for the SDT-based models and optimality. *Note.* Data points represent criterion values calculated over all trials of the last two sessions of each condition. In each panel, the main diagonal (dashed line) represents equality, the black solid line represents the linear regression, the numbers above panels denote values of the correlation coefficient (*r*) and its *p*-value, the regression slope (*b*_1_), and its intercept (*b*_0_). Note that the axis ranges for Model 2 had to be expanded and therefore are different from that of all other panels.

A satisfactory model should not only proffer a high positive correlation between predicted and obtained values, but also correct parameter estimates, and we can assess the precision of the predictions by the slopes and intercepts of the regression lines. While all regression lines had intercepts very close to 0 (ranging from –0.03 to 0.02), only Model 3 produced slopes which were not significantly different from 1 in both experiments (slopes were 0.95 (*p*=.79) and 1.12 (*p*=.56), for Experiments 1 and 2, respectively). The slopes of Model 1 and the optimality account in Experiment 1 were 0.67 and 0.61 and both were significantly smaller than 1 (*p*=.004 and *p*=.0004, respectively). Hence, the bad performance of Model 1 can indeed by attributed to its inability to explain the steady-state behavior of the animals. Unfortunately, the close correspondence between predicted and obtained criteria for Model 3 does not automatically validate it. Recall that fits of this model yielded negative values for the learning rate parameter υ for all four animals (Figure 4), and as argued above, a negative learning parameter not only violates common sense, but simulating Model 3 with negative υ also fails to generate behavior that is similar to that of the animals (Supplemental Figure 2). To sum up, none of the three SDT-based criterion learning models provides a satisfactory account of the experimental data.

### A trial-by-trial version of the Davison-Tustin model

While comparatively little research effort has been directed toward describing trial-by-trial changes of response bias in perceptual discrimination tasks, there exists an established framework for modeling steady-state response bias as a function of experimental parameters (reviewed in Commons et al., 1991). In a seminal article, Davison and Tustin (1978) proposed a behavioral model of signal detection based on the generalized matching law (Baum, 1974). Their model (henceforth, DT model) predicts that animals match response ratios to reinforcement ratios (as in the GLM) for each of the stimuli, and that the degree of matching is affected by stimulus discriminability (expressed as a sensitivity parameter log *d*), sensitivity to reinforcement (slope *a*), and a general bias parameter (intercept log *c*; see Methods for details). Unlike standard depictions of discrimination behavior, the DT model describes performance in a coordinate system formed by the logarithm of the reinforcement ratio (log Rf_R2_/Rf_R1_) on the abscissa and the logarithm of the response ratio (log R2/R1) on the ordinate. The DT model successfully describes behavioral performance in a wide range of different experiments (e.g. (Davison & McCarthy, 1980; McCarthy et al., 1982; McCarthy & Davison, 1980, 1984) and has since then been modified and extended to encompass multi-stimulus, multi-response procedures (e.g., Alsop, 1991; Davison, 1991; Davison & Jenkins, 1985; Davison & Nevin, 1999). Prompted by the failure of the three SDT-based models, we therefore turned to assess the fit of the DT model to steady-state response bias.

Figure 6 shows each animal’s steady-state response ratios (calculated over the last two sessions of each condition) as a function of the obtained reinforcement ratio in logarithmic coordinates for both Experiment 1 (left column) and Experiment 2 (right column). The DT model can be fitted easily through separate linear regressions for S1 and S2 trials. Most of the response data is reasonably well captured by linear fits, with the notable exception of S2 trials in Experiment 2 of subject AD1.

**Figure 6.**
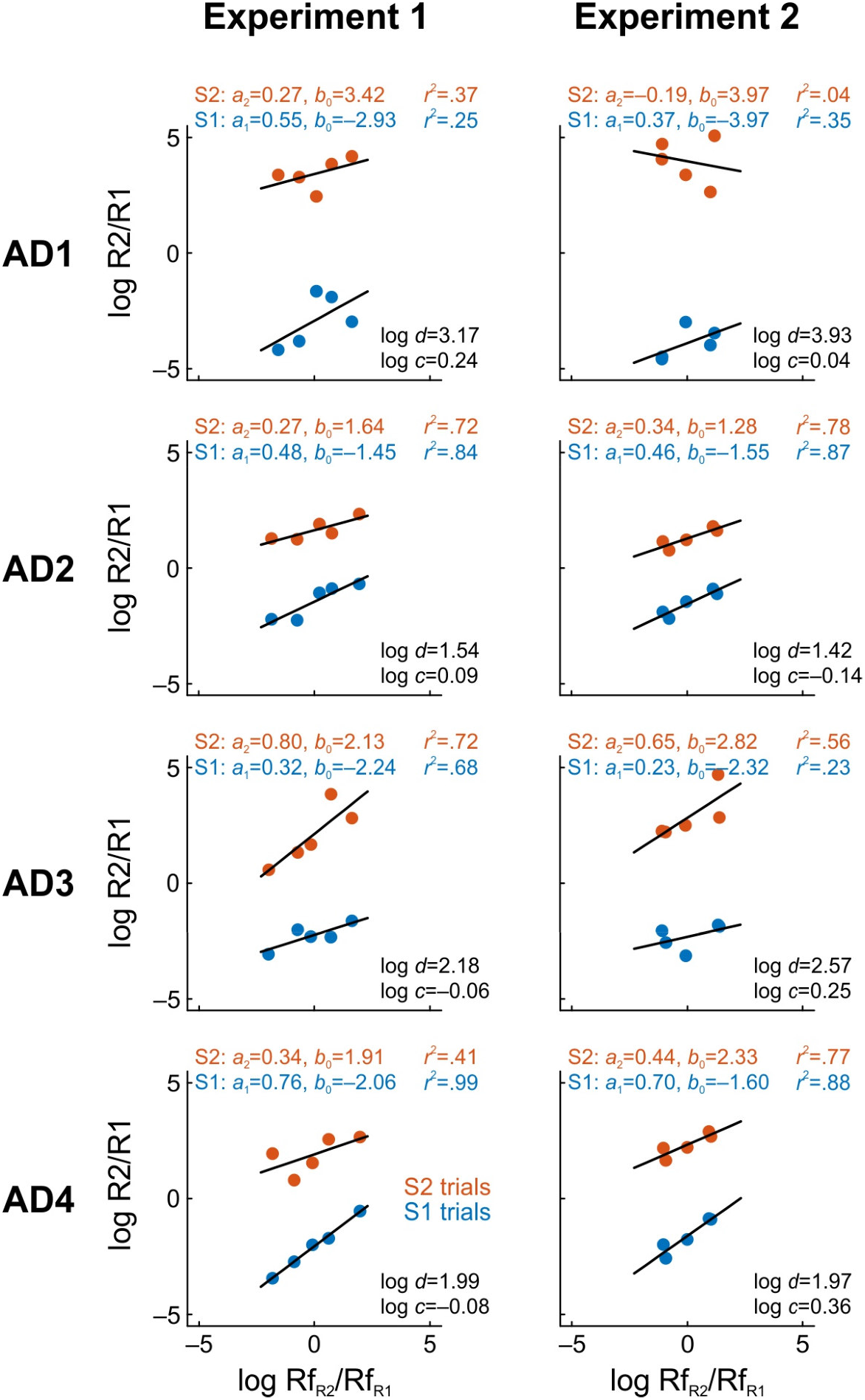
Fits of the Davison-Tustin model. *Note.* In all panels, data points show the logarithmic reinforcement ratios, log (Rf_R2_/Rf_R1_), plotted against the logarithm of the response ratios, log (R2/R1), computed over the total responses in the last two sessions of each condition and separately for S1 (blue) and S2 (red) trials (two lines per panel). Left column panels pertain to Experiment 1, right column panels to Experiment 2. Log *d* and log *c* are the parameters of the DT model, parameters *a*_1_ and *a*_2_ represent regression slopes in S1 and S2 trials, respectively, *b*_0_ represents the corresponding intercepts, and *r*^2^ is the squared correlation coefficient.

To the best of our knowledge, there exists no trial-by-trial instantiation of the DT model so far. We therefore developed such a trial-based version of the DT model (henceforth, tb-DT model) to investigate whether this provides a more satisfactory account of the data than the three SDT-based models. Our tb-DT model uses leaky integration of the history of past reinforcements to estimate the reinforcement ratio in each trial, the same mechanism employed by the SDT-based models (parameter γ). The choice of R1 over R2 in trial *t* is made probabilistically on the basis of the estimate of the reinforcement ratio (multiplied by *a*_x_, a parameter of sensitivity to reinforcement for stimulus *x*), the stimulus discriminability parameter log *d* and the stimulus-independent bias parameter log *c* (see Methods for details).

Figure 7A shows the maximum-likelihood fits of the tb-DT model to the data in the same format as Figure 3 did for the three SDT-based models. The parameter estimates for each animal are given in Table 3. Qualitatively, the tb-DT model provided a superior fit to the data than any of the three SDT-based models. BIC values of the tb-DT model (see Table 4) were considerably smaller than those of Model 2 (range 613–999) and also Model 1 for three of four animals (–47, 182, 301, 80 for subjects AD1 thru AD4, respectively). On the other hand, the BIC values were larger than those of Model 3 for three of four animals (–509, –127, 1, –311 for AD1 thru AD4, respectively). However, while models with low BIC values are usually preferred, a low BIC value by itself is not sufficient to endorse a model. In the present situation, Model 3 is to be rejected nonetheless, in part because forward simulations with negative learning rates generate behavior that is qualitatively very different from that of the subjects (Supplemental Figure 2), which however is not the case for simulations of the tb-DT model (Supplemental Figure 4).

**Figure 7.**
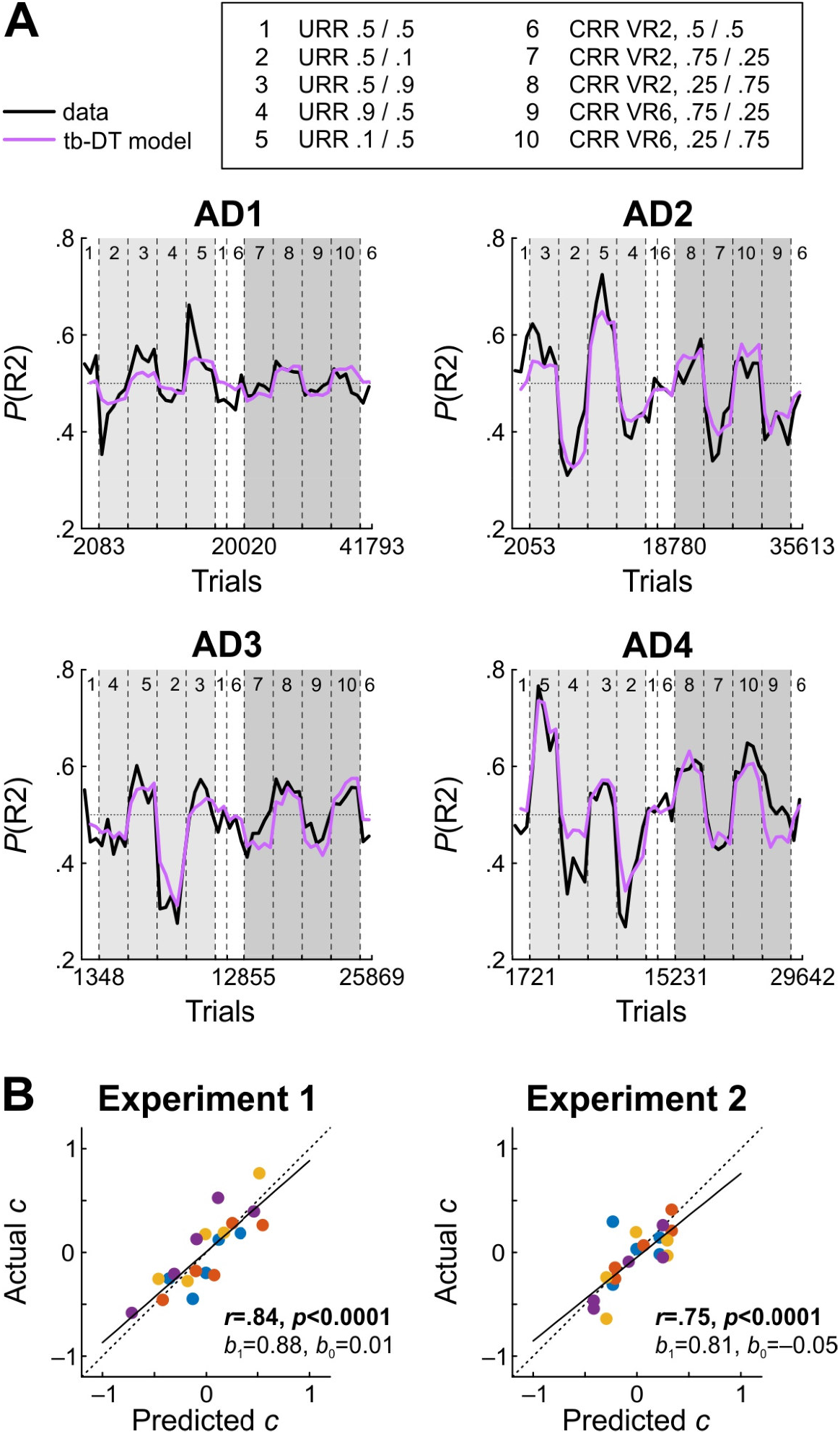
Maximum-likelihood fits of the trial-based version of the DT model and steady-state predictions. *Note.* A) Fits of the trial-based DT model to each subject’s data. Empirical *P*(R2) values redrawn in black from Figure 2. B) Predicted steady-state criterion values derived on the basis each subject’s fitted tb-DT model parameters. Conventions as in Figure 5.

**Table 3.** Fitted parameters of the trial-by-trial version of the DT model.

| Subject |  | Parameters |  |  |  |
| --- | --- | --- | --- | --- | --- |
| ID | $\log d$ | $\log c$ | $a_1$ | $a_2$ | $\gamma$ |
| AD1 | 3.186 | 0.01 | 0.542 | 0.378 | 0.983 |
| AD2 | 1.43 | -0.111 | 0.487 | 0.389 | 0.99 |
| AD3 | 2.233 | 0.018 | 0.35 | 0.697 | 0.988 |
| AD4 | 1.897 | 0.15 | 0.641 | 0.49 | 0.99 |

*Note.* The values of the leaky integration parameter $\gamma$ can be expressed as exponential half-times in723 trials, $\log .5 / \log \gamma$ . For subjects AD1 thru AD4, these correspond to 41, 70, 57, and 72 trials.

**Table 4.**
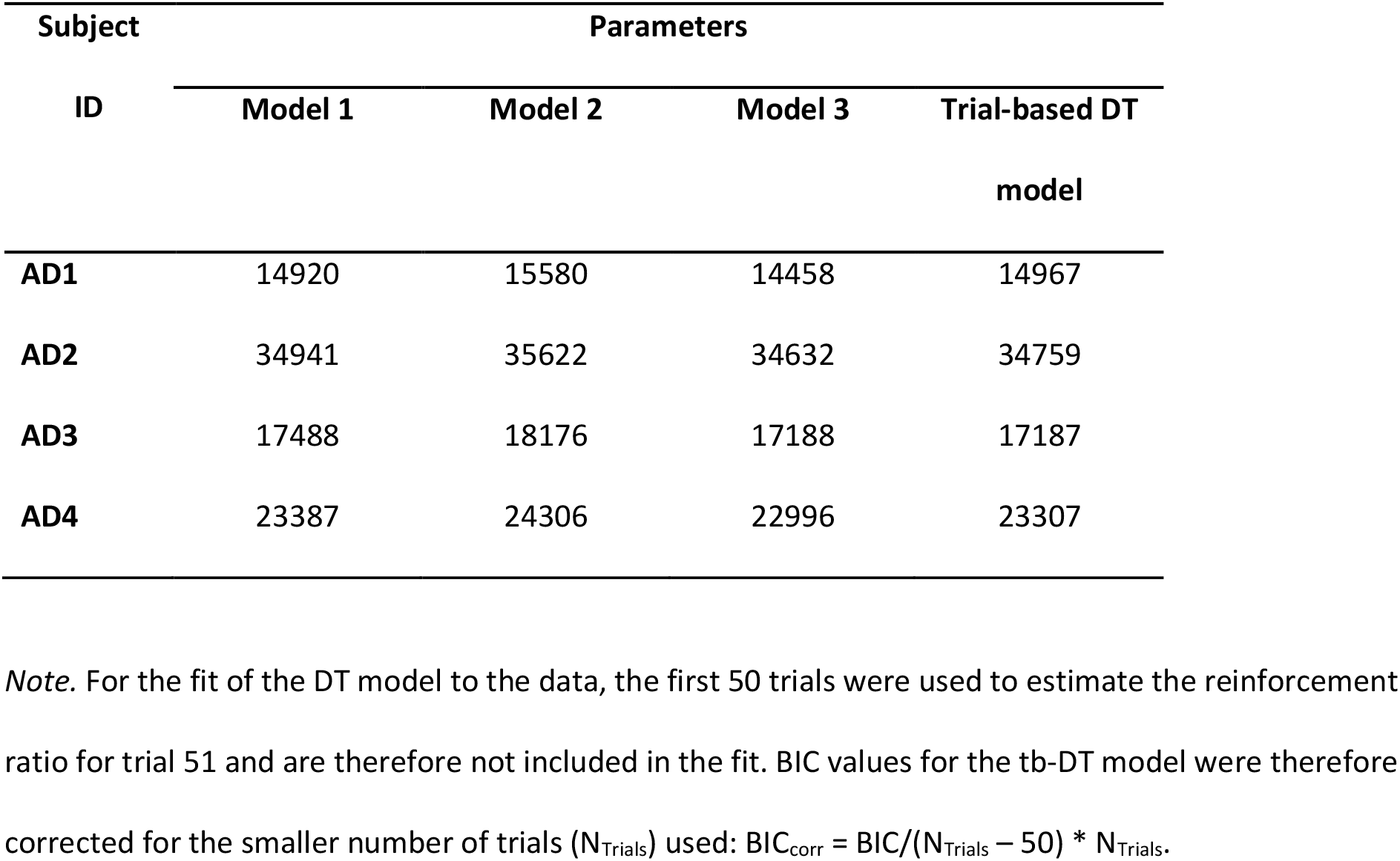
Bayesian Information Criterion (BIC) for all models.

Lastly, we examined the extent to which the steady-state predictions of the DT model are borne out by the data. The scatterplots in Figure 7B show close correspondence between the two, not only in terms of high and significant values of *r*, but also in terms of the slopes of the regression lines which were 0.88 and 0.81 for Experiments 1 and 2, respectively, and both slopes were not significantly different from 1 (*p*=.37 and *p*=.27).

## Discussion

While signal detection theory is considered by many to be perhaps “the most towering achievement of basic psychological research of the last half century” (Estes, 2002), relatively little work has been directed towards uncovering the mechanisms underlying criterion learning, and the attempts conducted so far have yielded incoherent results (compare e.g. Dorfman et al., 1975; Dorfman & Biderman, 1971; Hautus et al., 2022; Kac, 1962; Stüttgen et al., 2011, 2013; this study). In an effort to begin clarifying this issue, we chose to compare three simple models of criterion learning in their ability to fit experimental data. Because none of the three models provided a satisfactory fit to the data, we additionally derived a dynamic version of behavioral detection model (Davison & Tustin, 1978).

### Does any of the three criterion learning models proffer a satisfactory description of criterion learning?

The income-based Model 1, which updates the criterion on reinforced trials only, provided a reasonable fit, although the fitted choice probabilities differed from the data considerably especially in conditions 2 and 5 (Figure 3). Also, steady-state predicted criterion values were often considerably smaller than those observed (Figure 5). Forward simulations showed that this model predicts more extreme criteria in conditions 3 and 4 than conditions 2 and 5, respectively (Supplemental Figure 2), which was however observed only in two out of four animals. For Experiment 2, forward simulations predict that conditions 7 and 8 (with reinforcement delivered at VR 2) produce more extreme criterion values than conditions 9 and 10 (VR 6), which implies that reinforcement density (i.e., number of expected reinforcers per trial) affects criterion placement. However, this prediction was again met in two animals only.

On the other hand, fits of Model 2, which updates the criterion after non-reinforced responses only, largely failed to account for the data (Figure 3). Forward simulations of this model demonstrated its inability to produce systematic criterion shifts when supplied with a negative learning rate parameter (Supplemental Figure 2), and its steady-state predictions aligned badly with observed criteria (Figure 5).

Model 2 learns after all non-reinforced responses, which in our experiments include a large number of correct responses. Accordingly, one might ask whether animals might learn on error trials only. Learning from errors is considered to be a good description of human criterion setting performance (Dorfman, 1969; Dorfman et al., 1975; Friedman et al., 1968; Kac, 1962; Killeen et al., 2018; Thomas, 1975). However, in our experiments, an agent purely learning from errors would not be affected by asymmetric reinforcement probabilities *at all* and would therefore produce no shifts in response to changing reinforcement contingencies (see Supplemental Figure 2 for forward simulations), which is at odds with the systematic biases we observed. Moreover, fitting a pure error-learning model to the data produced negative learning rates for all four animals (data not shown).

Model 3, which learns after both reinforced and non-reinforced responses, had by far the lowest BIC values and provided an excellent fit to the data of all animals, often capturing even minor session-to-session fluctuations (Figure 3). Unfortunately, this model still cannot be considered further, because the fitted learning rate parameter υ turned out negative for all animals. This has the effect that, after an error, the criterion is shifted such that the same error becomes more likely to occur on the next trial. This does not only make no sense from a theoretical point of view; in addition, Model 3 with negative υ failed badly to recapture the observed criterion setting behavior when simulated forward (see Supplemental Figure 2 and Corrado et al., 2005, for a similar case). The reason Model 3 fits the data so much better than its competitor, Model 1, is likely because it exploits lose-stay patterns in the experimental data which arise as a consequence of response bias.

In conclusion, none of the three criterion learning models provided a satisfactory account of the behavioral data. Although Model 1 was able to capture and generate criterion shifts in the predicted direction, there is a lot of variance left unaccounted for by this model. Models 2 and 3, on the other hand, could be clearly rejected as viable mechanistic accounts of criterion learning.

### Does the DT model proffer an adequate description of criterion learning?

In animal psychophysics, subjects usually work for food or water reinforcement. Thus, from the perspective of the animal, there is no fundamental difference between a perceptual decision-making task and other reinforcement-based choice procedures. Accordingly, it is straightforward to connect SDT to the most important description of reinforcement-based choice behavior, the (generalized) matching law (Baum, 1974; Herrnstein, 1961). This was done in classic work by Davison and Tustin (1978) and then developed further (Alsop, 1991; Davison, 1991; Davison & Jenkins, 1985; Davison & Nevin, 1999).

The DT model has been successful in fitting data from a diverse set of experiments (Davison & McCarthy, 1980; McCarthy & Davison, 1979, 1980, 1981, 1984), including our data (Figure 6). Moreover, it predicts the steady-state criterion values better than any of the three criterion learning models (compare Figure 5 and Figure 7A), with the exception of Model 3 which however is untenable for other reasons. One major difference between the criterion learning models and the DT model is how reinforcers are assumed to determine response bias. The learning models conceptualize criterion placement as resulting from the *difference* in the number of reinforcers obtained from the two responses (scaled by reinforcement density, i.e. the overall frequency of reinforcement, because the criterion drifts back to 0 on trials without reinforcement). In contrast, the DT model posits that response bias results from the *ratio* of reinforcers obtained from the two responses. Incidentally, the DT model does not assume any effect of reinforcement density. While we did not explicitly test whether reinforcement density has an effect, the highly similar steady-state criterion values for VR 2 and VR 6 in Experiment 2 are consistent with this prediction.

To our knowledge, this is the first study to propose and examine a trial-based version of the DT model. Although the original model has been modified and extended considerably (Alsop, 1991; Davison, 1991; Davison & Jenkins, 1985; Davison & Nevin, 1999), we chose to build on the classic DT model for several reasons. First, the DT model is considerably easier to fit than its successor models, and the same holds for the trial-based version. Second, both for our as for other two-stimulus, two-response data sets, the DT model already provides a very good fit (Figures 6 and 7A; and e.g. McCarthy & Davison, 1979, 1980). Third, simulations of the latest version of the model (Davison & Nevin, 1999) yielded results which were very similar to those shown in Supplemental Figure 4.

The tb-DT model provided a superior fit for three of the four subjects compared to the SDT-based criterion learning models. However, it clearly failed to capture the behavior of subject AD1 in Experiment 1. Conspicuously, this subject had by far the highest values of *d’* and log *d* and exhibited prominent and transient extreme values of *P*(R2) in conditions 2 and 5 which featured the most extreme reinforcer ratios. Similar “overshooting” behavior was seen in an earlier study with pigeons (Stüttgen, Yildiz, et al., 2011). Also, simulations of the tb-DT model with parameters similar to those of our subjects were qualitatively different from observed behavior (Supplemental Figure 4); in particular, adaptation to novel conditions was considerably more sluggish, and this did not change much when simulations were run with lower values of γ.

At a more general level, the DT model has several important limitations which also apply to its trial-by-trial instantiation. First, the sensitivity-to-reinforcement parameter *a* lacks theoretical justification and has repeatedly been found not to be independent from the discriminability parameter log *d* (e.g., Davison & Jenkins, 1985). Second, it does not explain how subjects come to be biased by the stimuli in the first place, i.e., how they learn to associate S1 with R1 and S2 with R2 (this is much clearer in successor models, see e.g. (Davison & Nevin, 1999). Third, from the point of view of signal detection models, the animals do not know whether they are in an S1 or S2 trial, but the tb-DT model equations use this information to compute the animals’ response probabilities. Fourth, the DT model is limited to two-stimulus, two-response procedures (although successors to the DT model are not, see Davison, 1991, and our trial-by-trial instantiation is further restricted to discrete-trial procedures. Fifth, it is not clear how the scope of the DT model could be extended to encompass lapses (Pisupati et al., 2021), reaction times (Hernández-Navarro et al., 2021) or decision confidence (Kepecs et al., 2008; Lak, Hueske, et al., 2020). In the future, some of these limitations may be overcome by equipping an SDT-based criterion learning model with an updating mechanism based on the DT model or its successors.

## References

1. Alsop, B. (1991). Behavioral models of signal detection and detection models of choice. In M. L. Commons, J. A. Nevin, & M. C. Davison (Eds.), Signal detection: Mechanisms, models, and applications (pp. 39–55). Erlbaum. 10.4324/9780203772430

2. Alsop, B. (1998). Receiver operating characteristics from nonhuman animals: Some implications and directions for research with humans. Psychonomic Bulletin & Review, 5(2), 239–252. 10.3758/BF03212946

3. Baum, W. M. (1974). On two types of deviation from the matching law: bias and undermatching. Journal of the Experimental Analysis of Behavior, 22(1), 231–242. 10.1901/jeab.1974.22-231

4. Benjamin, A. S., Diaz, M., & Wee, S. (2009). Signal detection with criterion noise: applications to recognition memory. Psychological Review, 116(1), 84–115. 10.1037/a0014351

5. Boneau, C. A., & Cole, J. L. (1967). Decision theory, the pigeon, and the psychophysical function. Psychological Review, 74(2), 123–135. 10.1037/h0024287

6. Busemeyer, J. R., & Myung, I. J. (1992). An Adaptive Approach to Human Decision Making: Learning Theory, Decision Theory, and Human Performance. Journal of Experimental Psychology: General, 121(2), 177–194. 10.1037/0096-3445.121.2.177

7. Commons, M. L., Nevin, J. A., & Davison, M. C. (Eds.). (1991). Signal detection: Mechanisms, models, and applications. Erlbaum. 10.4324/9780203772430

8. Corrado, G. S., Sugrue, L. P., Sebastian Seung, H., & Newsome, W. T. (2005). Linear-Nonlinear-Poisson Models of Primate Choice Dynamics. Journal of the Experimental Analysis of Behavior, 84(3), 581– 617. 10.1901/jeab.2005.23-05

9. David M. Green, & John A. Swets. (1988). Signal Detection Theory and Psychophysics. Peninsula Publishing.

10. Davison, M. (1991). Stimulus discriminability, contingency discriminability, and complex stimulus control. In M. L. Commons, J. A. Nevin, & M. C. Davison (Eds.), Signal detection: Mechanisms, models, and applications (pp. 57–78). Erlbaum. 10.4324/9780203772430

11. Davison, M., & Jenkins, P. E. (1985). Stimulus discriminability, contingency discriminability, and schedule performance. Animal Learning & Behavior, 13(1), 77–84. 10.3758/BF03213368

12. Davison, M., & McCarthy, D. (1980). Reinforcement for Errors in A Signal-Detection Procedure. Journal of the Experimental Analysis of Behavior, 34(1), 35–47. 10.1901/jeab.1980.34-35

13. Davison, M., & McCarthy, D. (1981). Undermatching and structural relations. Behaviour Analysis Letters, 1(1), 67–72.

14. Davison, M., & Nevin, J. (1999). Stimuli, reinforcers, and behavior: an integration. Journal of the Experimental Analysis of Behavior, 71(3), 439–482. 10.1901/jeab.1999.71-439

15. Davison, M., & Tustin, R. D. (1978). The relation between the generalized matching law and signal-detection theory. Journal of the Experimental Analysis of Behavior, 29(2), 331–336. 10.1901/jeab.1978.29-331

16. Dorfman, D. D. (1969). Probability matching in signal detection. Psychonomic Science, 17(2), 103–103. 10.3758/BF03336468

17. Dorfman, D. D. (1973). Likelihood Function of Additive Learning Models – Sufficient Conditions for Strict Log-Concavity and Uniqueness of Maximum. Journal of Mathematical Psychology, 10(1), 73–85. 10.1016/0022-2496(73)90005-9

18. Dorfman, D. D., & Biderman, M. (1971). A learning model for a continuum of sensory states. Journal of Mathematical Psychology, 8(2), 264–284. 10.1016/0022-2496(71)90017-4

19. Dorfman, D. D., Saslow, C. F., & Simpson, J. C. (1975). Learning Models for A Continuum of Sensory States Reexamined. Journal of Mathematical Psychology, 12(2), 178–211. 10.1016/0022-2496(75)90056-5

20. Erev, I. (1998). Signal detection by human observers: a cutoff reinforcement learning model of categorization decisions under uncertainty. Psychological Review, 105(2), 280–298. 10.1037/0033-295X.105.2.280

21. Estes, W. K. (2002). Traps in the route to models of memory and decision. Psychonomic Bulletin and Review, 9(1), 3–25. 10.3758/BF03196254

22. Feng, S., Holmes, P., Rorie, A., & Newsome, W. T. (2009). Can monkeys choose optimally when faced with noisy stimuli and unequal rewards? PLoS Computational Biology, 5(2), e1000284. 10.1371/journal.pcbi.1000284

23. Friedman, M. P., Carterette, E. C., Nakatani, L., & Ahumada, A. (1968). Comparisons of some learning models for response bias in signal detection. Perception & Psychophysics, 3, 5–11. 10.3758/BF03212703

24. Funamizu, A. (2021). Integration of sensory evidence and reward expectation in mouse perceptual decision-making task with various sensory uncertainties. IScience, 24, 102826. 10.1016/j.isci.2021.102826

25. Gold, J. I., & Ding, L. (2013). How mechanisms of perceptual decision-making affect the psychometric function. Progress in Neurobiology, 103, 98–114. 10.1016/j.pneurobio.2012.05.008

26. Harley, C. B. (1981). Learning the evolutionarily stable strategy. Journal of Theoretical Biology, 89(4), 611–633. 10.1016/0022-5193(81)90032-1

27. Hautus, M. J., Macmillan, N. A., & Creelman, C. D. (2022). *Detection theory: a user’s guide* (3rd ed.). Routledge. 10.4324/9781003203636

28. Hernández-Navarro, L., Hermoso-Mendizabal, A., Duque, D., de la Rocha, J., & Hyafil, A. (2021). Proactive and reactive accumulation-to-bound processes compete during perceptual decisions. Nature Communications, 12(1). 10.1038/s41467-021-27302-8

29. Herrnstein, R. J. (1961). Relative and absolute strength of response as a function of frequency of reinforcement. Journal of the Experimental Analysis of Behavior, 4, 267–272. 10.1901/jeab.1961.4-267

30. Kac, M. (1962). A note on learning signal detection. IRE Transactions on Information Theory, 8(2), 126–128. 10.1109/TIT.1962.1057687

31. Kass, R. E., & Raftery, A. E. (1995). Bayes factors. Journal of the American Statistical Association, 90(430), 773–795. 10.1080/01621459.1995.10476572

32. Kepecs, A., Uchida, N., Zariwala, H. a, & Mainen, Z. F. (2008). Neural correlates, computation and behavioural impact of decision confidence. Nature, 455(7210), 227–231. 10.1038/nature07200

33. Killeen, P. R., Taylor, T. J., & Treviño, M. (2018). Subjects adjust criterion on errors in perceptual decision tasks. Psychological Review, 125(1), 117–130. 10.1037/rev0000056

34. Lak, A., Hueske, E., Hirokawa, J., Masset, P., Ott, T., Urai, A. E., Donner, T. H., Carandini, M., Tonegawa, S., Uchida, N., & Kepecs, A. (2020). Reinforcement biases subsequent perceptual decisions when confidence is low: A widespread behavioral phenomenon. ELife, 9, 1–26. 10.7554/eLife.49834

35. Lak, A., Nomoto, K., Keramati, M., Sakagami, M., & Kepecs, A. (2017). Midbrain Dopamine Neurons Signal Belief in Choice Accuracy during a Perceptual Decision. Current Biology, 27(6), 821–832. 10.1016/j.cub.2017.02.026

36. Lak, A., Okun, M., Moss, M. M., Gurnani, H., Farrell, K., Wells, M. J., Reddy, C. B., Kepecs, A., Harris, K. D., & Carandini, M. (2020). Dopaminergic and Prefrontal Basis of Learning from Sensory Confidence and Reward Value. Neuron, 105(4), 700–711. 10.1016/j.neuron.2019.11.018

37. Luce, R. D. (1963). A Threshold Theory for Simple Detection Experiments. Psychological Review, 70(1), 61–79. 10.1037/h0039723

38. McCarthy, D., & Davison, M. (1979). Signal Probability, Reinforcement and Signal-Detection. Journal of the Experimental Analysis of Behavior, 32(3), 373–386. 10.1901/jeab.1979.32-373

39. McCarthy, D., & Davison, M. (1980). Independence of Sensitivity to Relative Reinforcement Rate and Discriminability in Signal-Detection. Journal of the Experimental Analysis of Behavior, 34(3), 273–284. 10.1901/jeab.1980.34-273

40. McCarthy, D., & Davison, M. (1981). Towards a behavioral theory of bias in signal detection. Perception & Psychophysics, 29(4), 371–382. 10.3758/bf03207347

41. McCarthy, D., & Davison, M. (1984). Isobias and alloiobias functions in animal psycophysics. Journal of Experimental Psychology: Animal Behavior Processes, 10(3), 390–409. 10.1037/0097-7403.10.3.390

42. McCarthy, D., Davison, M., & Jenkins, P. E. (1982). Stimulus discriminability in free-operant and discrete-trial detection procedures. Journal of the Experimental Analysis of Behavior, 37(2), 199–215. 10.1901/jeab.1982.37-199

43. Mill, R. W., Alves-Pinto, A., & Sumner, C. J. (2014). Decision Criterion Dynamics in Animals Performing an Auditory Detection Task. PLoS ONE, 9(12), e114076. 10.1371/journal.pone.0114076

44. Pisupati, S., Chartarifsky-Lynn, L., Khanal, A., & Churchland, A. K. (2021). Lapses in perceptual decisions reflect exploration. ELife, 10, 1–27. 10.7554/ELIFE.55490

45. Stoilova, V. V., Knauer, B., Berg, S., Rieber, E., Jäkel, F., & Stüttgen, M. C. (2020). Auditory cortex reflects goal-directed movement but is not necessary for behavioral adaptation in sound-cued reward tracking. Journal of Neurophysiology, 124(4), 1056–1071. 10.1152/jn.00736.2019

46. Stubbs, D. A., & Pliskoff, S. S. (1969). Concurrent Responding with Fixed Relative Rate of Reinforcement. Journal of the Experimental Analysis of Behavior, 12(6), 887-. 10.1901/jeab.1969.12-887

47. Stüttgen, M. C., Kasties, N., Lengersdorf, D., Starosta, S., Güntürkün, O., & Jäkel, F. (2013). Suboptimal criterion setting in a perceptual choice task with asymmetric reinforcement. Behavioural Processes, 96, 59–70. 10.1016/j.beproc.2013.02.014

48. Stüttgen, M. C., Schwarz, C., & Jäkel, F. (2011). Mapping spikes to sensations. Frontiers in Neuroscience, 5(November), 125.

49. Stüttgen, M. C., Yildiz, A., & Güntürkün, O. (2011). Adaptive criterion setting in perceptual decision making. Journal of the Experimental Analysis of Behavior, 96(2), 155–176. 10.1901/jeab.2011.96-155

50. Swets, J. A. (1961a). Detection theory and psychophysics: A review. Psychometrika, 26(1), 49–63. 10.1007/BF02289684

51. Swets, J. A. (1961b). Is there a sensory threshold? Science, 134(3473), 168–177. 10.1126/science.134.3473.168

52. Teichert, T., & Ferrera, V. P. (2010). Suboptimal integration of reward magnitude and prior reward likelihood in categorical decisions by monkeys. Frontiers in Neuroscience, 4(November), 186. 10.3389/fnins.2010.00186

53. Thomas, E. A. C. (1975). Criterion adjustment and probability matching. Perception & Psychophysics, 18, 158–162. 10.3758/BF03204104

54. Treisman, M., & Williams, T. C. (1984). A theory of criterion setting with an application to sequential dependencies. Psychological Review, 91(1), 68–111. 10.1037//0033-295X.91.1.68

55. Vandevelde, J. R., Yang, J.-W., Albrecht, S., Lam, H., Kaufmann, P., Luhmann, H. J., & Stüttgen, M. C. (2023). Layer– and cell-type-specific differences in neural activity in mouse barrel cortex during a whisker detection task. Cerebral Cortex, 33, 1361–1382. 10.1093/cercor/bhac141

56. Wichmann, F. A., & Jäkel, F. (2018). Methods in Psychophysics. In J. T. Wixted (Ed.), Stevens’ Handbook of Experimental Psychology and Cognitive Neuroscience (4th ed., pp. 1–42). John Wiley & Sons, Inc. 10.1002/9781119170174.epcn507

57. Yang, T., & Shadlen, M. N. (2007). Probabilistic reasoning by neurons. Nature, 447(7148), 1075–1080. 10.1038/nature05852

