## Supplemental Figures for "Influence of Reinforcement and Its Omission on Trial-by-Trial Changes of Response Bias in Perceptual Decision-Making"

**Supplemental Figure 1**

*Behavioral paradigm and results overview for Experiments 1 and 2*

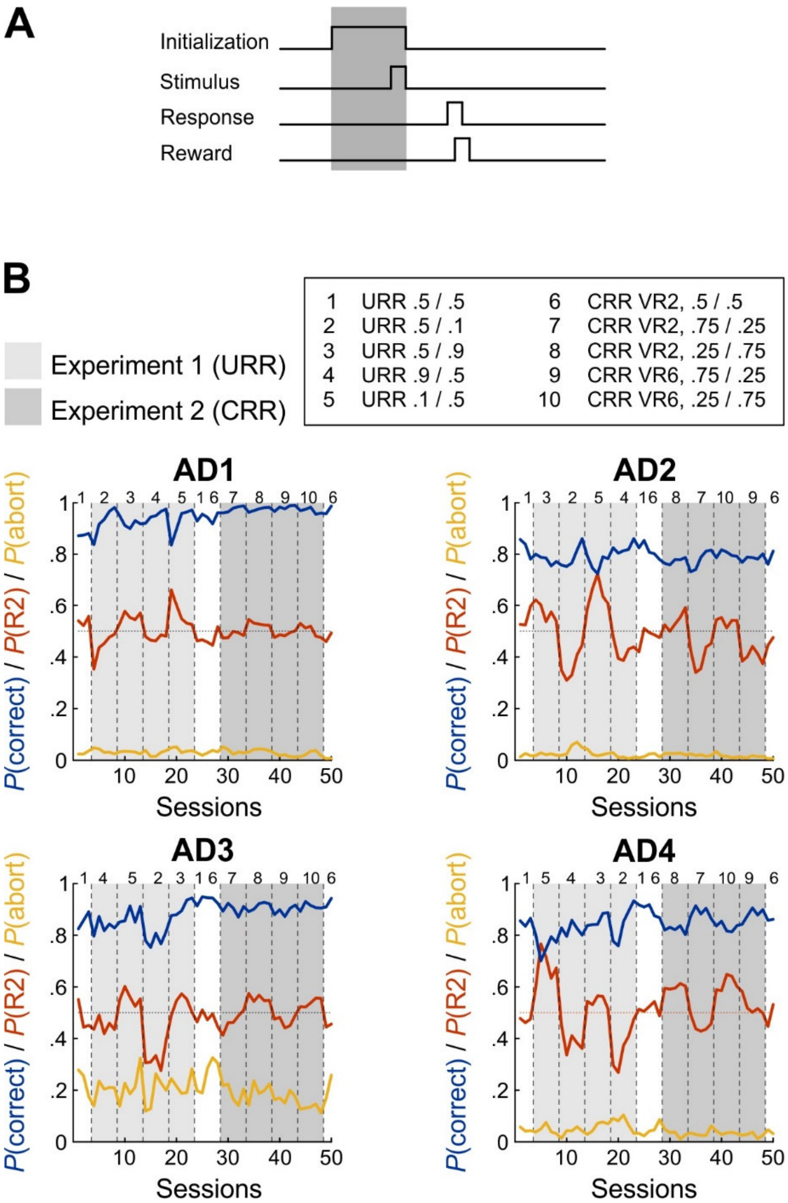


*Note.* A) Schematic outline of epochs in a single trial of the behavioral task. Gray shaded area denotes time interval in which rats had to maintain nose poking at the center port. B) Proportion of correct trials, *P*(correct) proportion of R2, *P*(R2), and proportion of aborted trials, *P*(abort), for all four subjects (AD1 thru AD4) for both experiments. All conventions as in Figure 2.

**Supplemental Figure 2**

*Example simulations of Models 1, 2 and 3 and an error-based learning model for both experiments*

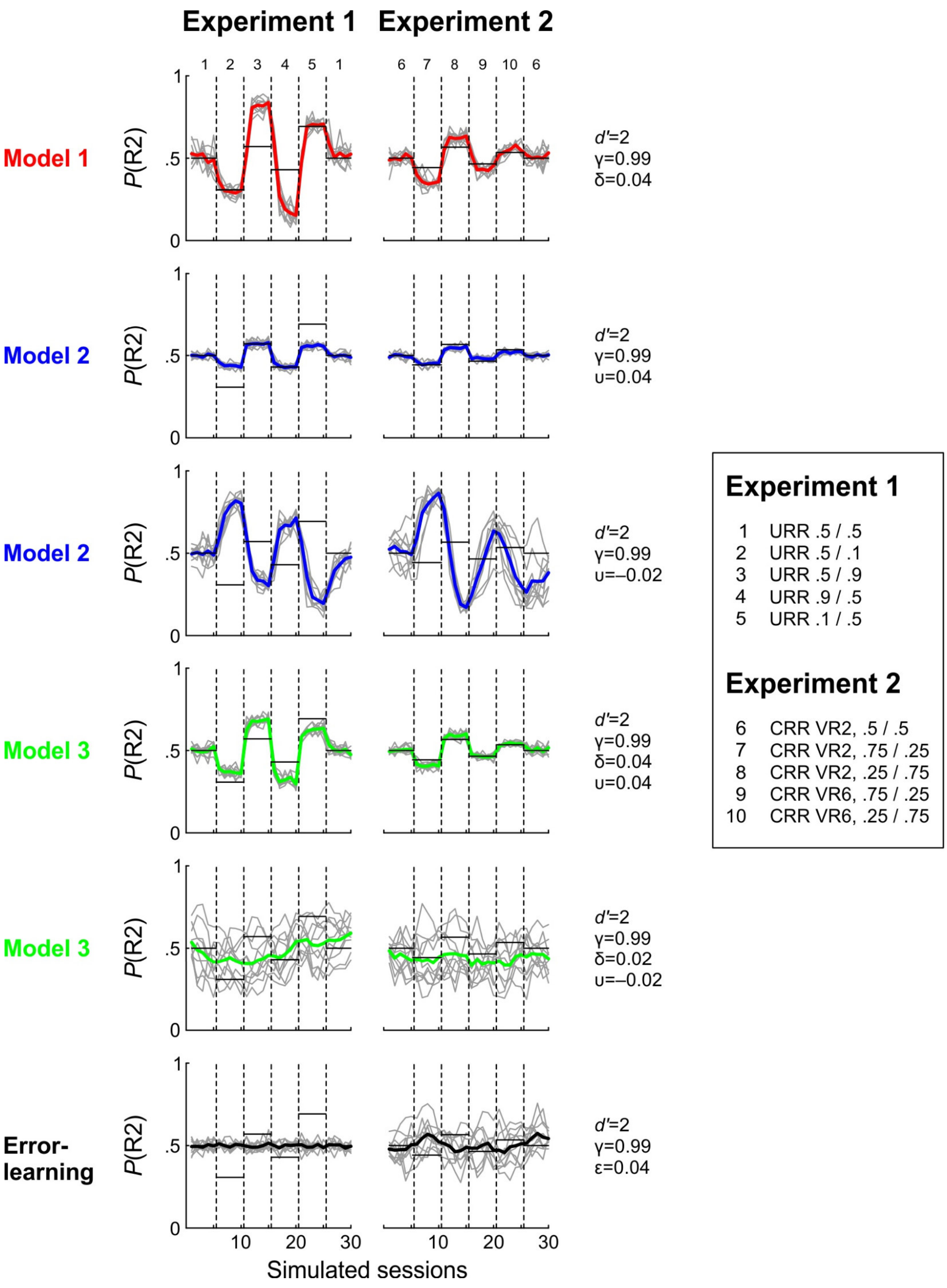

*Note.* Each model was simulated forward 10 times with random stimulus sequences. Each condition consisted of 1500 trials which were split up into blocks of 6 virtual sessions with 300 trials each. In each panel, the 10 individual simulated sessions are shown as thin gray lines, their averages as bold lines. Dashed vertical lines separate the conditions, horizontal solid lines give the position of the optimal bias for reference. Model parameters are indicated on the right of each panel. Models 2 and 3 were run twice, once with positive values for both δ and υ, and once with a negative value of υ (as was found for the fits to the animals' data). Here, learning rate parameters δ and υ were reduced to 0.02 and –0.02, respectively, because values of 0.04 and –0.04 consistently produced exclusive choice in all simulations. Additionally, a pure error-learning model (Kac, 1962) was simulated which was identical to Model 2 with the exception that only incorrect rather than all trials without reinforcement led to a criterion shift with magnitude ε. Optimal *P*(R2) values for Experiment 2 were obtained through numerical optimization over $10^7$ trials.

**Supplemental Figure 3**

*Predictions of criterion location for optimal performance as well as models 1 and 2*

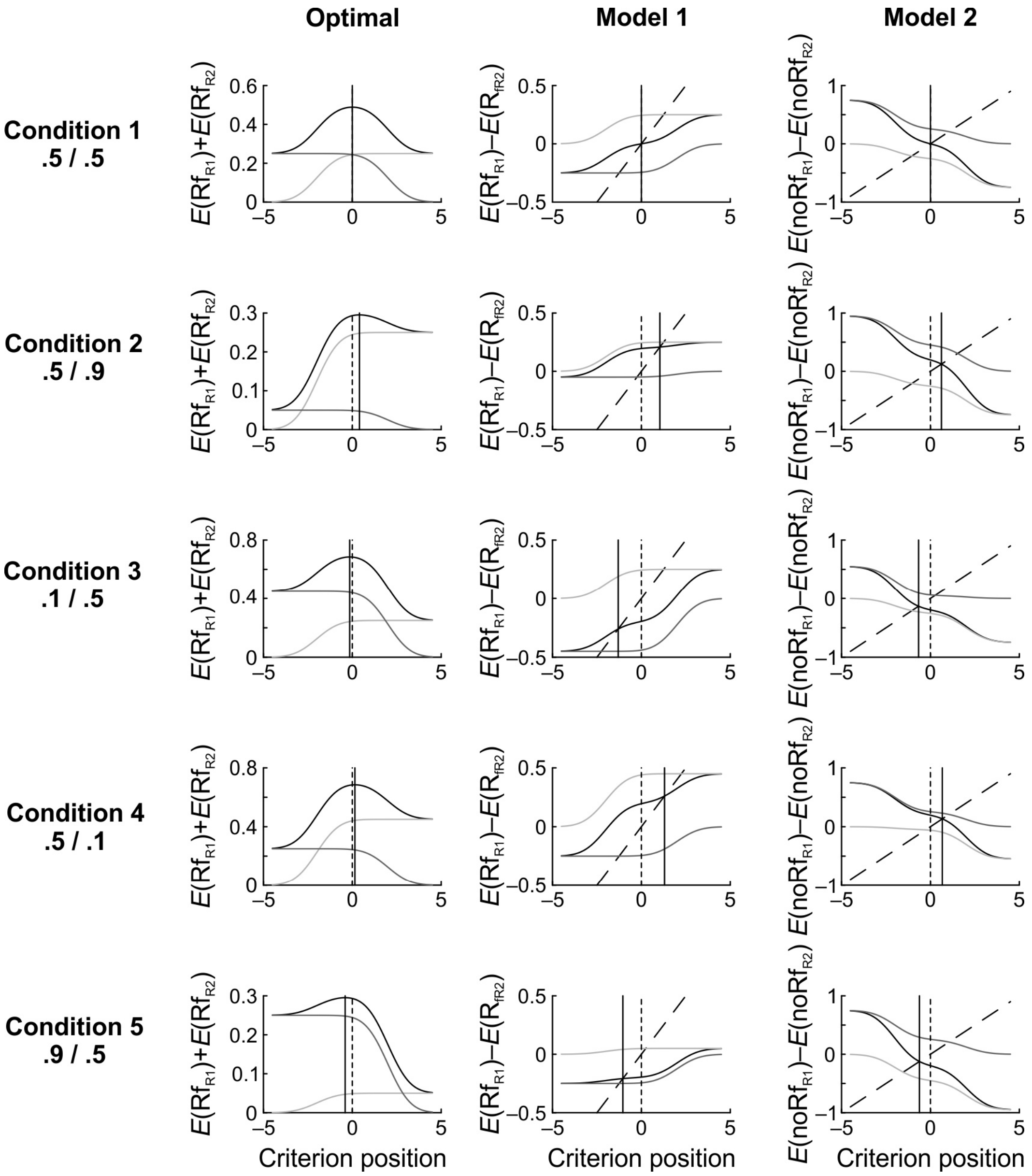


*Note.* In all plots, *d′*=2 and γ=0.99. For Models 1 and 2, δ=υ=0.04. In the left column, criterion predictions for optimal performance are shown. The bold black line is the objective reward function (ORF), which represents the total expected probability of reinforcement in a trial dependent on the criterion. The gray lines are the probabilities for reinforcement following each of the responses respectively. Optimal performance is achieved at the maximum of the ORF; the corresponding criterion is plotted as a thin black line. For reference, the neutral criterion at zero is shown as a dotted black line. In the middle and right column, criterion predictions for Models 1 and 2 respectively are shown. The difference between the expected probabilities for a trial with/without reinforcement following each response is plotted as a bold black line. The grey lines are these expected probabilities for each response respectively. Additionally, a straight line through zero is plotted as a dashed black line, whose slope depends on the leakage term $\gamma$ and the step size $\delta$ or $\upsilon$: it is $(1–\gamma)/\delta$ for model 1 and $(1–\gamma)/\upsilon$ for model 2. The predicted criterion location for the models is at the intersection of this straight line with the bold black line. For reference, the neutral criterion at zero is shown as a dotted black line.

**Supplemental Figure 4**

*Example simulations of the trial-based version of the DT model for both experiments*

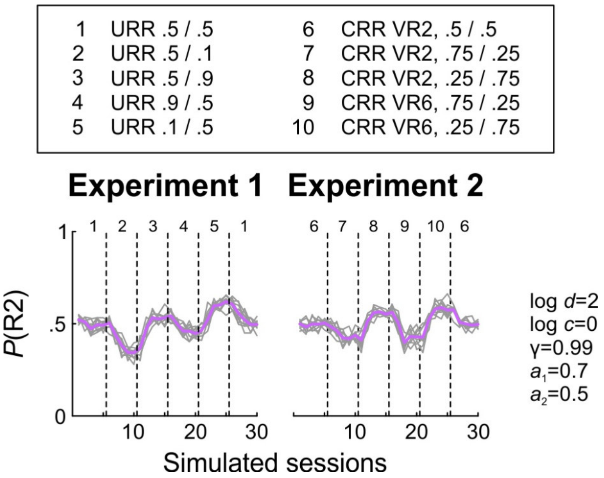


*Note.* Model simulations were conducted as described in Supplemental Figure 2, and the same conventions apply in this figure.
